# One-shot identification of SARS-CoV-2 S RBD escape mutants using yeast screening

**DOI:** 10.1101/2021.03.15.435309

**Authors:** Irene Francino Urdaniz, Paul J. Steiner, Monica B. Kirby, Fangzhu Zhao, Cyrus M. Haas, Shawn Barman, Emily R. Rhodes, Linghang Peng, Kayla G. Sprenger, Joseph G. Jardine, Timothy A. Whitehead

**Affiliations:** Department of Chemical and Biological Engineering, University of Colorado, Boulder, CO 80305, USA; The Scripps Research Institute, La Jolla, CA, USA; International AIDS Vaccine Initiative, New York, NY, USA

## Abstract

The potential emergence of SARS-CoV-2 Spike (S) escape mutants is a threat to reduce the efficacy of existing vaccines and neutralizing antibody (nAb) therapies. An understanding of the antibody/S escape mutations landscape is urgently needed to preemptively address this threat. Here we describe a rapid method to identify escape mutants for nAbs targeting the S receptor binding site. We identified escape mutants for five nAbs, including three from the public germline class VH3-53 elicited by natural COVID-19 infection. Escape mutations predominantly mapped to the periphery of the ACE2 recognition site on the RBD with K417, D420, Y421, F486, and Q493 as notable hotspots. We provide libraries, methods, and software as an openly available community resource to accelerate new therapeutic strategies against SARS-CoV-2.

**One Sentence Summary:** We present a facile method to identify antibody escape mutants on SARS-CoV-2 S RBD.

## MAIN

The type I viral fusion protein Spike (S) is a major antigenic determinant of SARS-CoV-2 and is the antigen used in all approved COVID-19 vaccines *(1–3)*. Recently, the B.1.1.7 (N501Y; U.K.), B.1.351 (E484K; South Africa), B.1.427 (L452R; California), and B.1.526 (S477N, E484K; NY) viral lineages have emerged. All encode single nucleotide substitutions in the S receptor binding domain (RBD) near recognition site for its cellular target angiotensin-converting enzyme 2 (ACE2) *(3–5)*.

Dozens of studies have reported the structural, epitopic, and functional landscape of non-neutralizing monoclonal antibodies and nAbs targeting trimeric S *(6–8)*. A prophetic understanding of the mutations on S that could evade antibody recognition would enable development of better vaccine boosters and monoclonal antibody therapies. Thus, we sought to develop an S RBD yeast surface display (YSD) platform (**Fig. S1**) *(9)*, as we hypothesized that broad identification of SARS-CoV-2 S escape mutants could be found by integrating high throughput screening platforms with deep sequencing. While a similar platform uses the loss of nAb binding to identify escape mutants *(10, 11)*, we rationalized that a functional screening assay that directly measures the ability of a nAb to compete with ACE2 for S RBD binding, would be a comparatively strong predictor of RBD escapability, as it accounts for mutations in RBD that would disrupt S binding to ACE2.

We had previously developed an aglycosylated S-RBD YSD platform (S RBD(333-537)-N343Q) *(8)* that can bind specifically to ACE2 (**Fig. 1a**). This S RBD construct has its one native N-linked glycan removed (N343Q) as the heavy N-linked mannosylation endemic of *S. cerevisiae* could hamper anti-S RBD mAb recognition. Cell surface titrations of CR3022 IgG and nAb HKU-910-30 IgG yielded apparent dissociation constants comparable to reported *in vitro* results *(6,8)* (**Fig. S2**). We next tested a panel of eleven additional anti-S RBD mAbs for binding to aglycosylated RBD *(7)*. Ten of the eleven mAbs recognized aglycosylated S RBD (**Fig. 1b**). The one panel member that did not bind, CC6.33, selectively recognizes the S309 epitope on the RBD containing the N-linked glycan at position 343 *(12)*.

**Figure 1.**
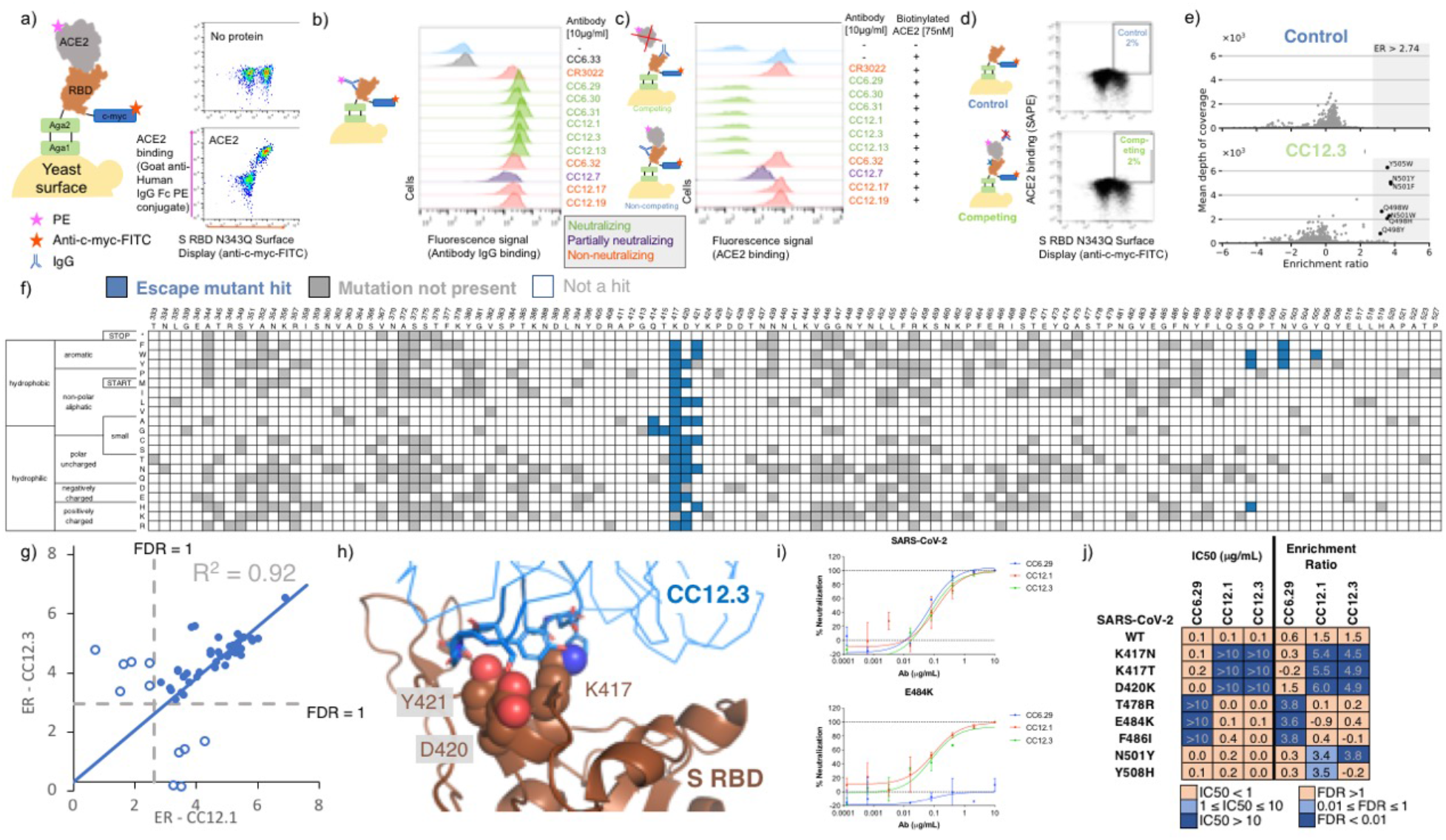
Identification and validation of SARS-CoV-2 S RBD escape mutants using yeast screening. **a)** Cartoon of the yeast display construct S-RBD(333-537)-N343Q. Cytograms show specific binding in the presence, but not absence, of ACE2-Fc. **b)** Binding profiles of aglycosylated S RBD labeled with 10 μg/mL of indicated mAb. Antibodies are color coded according to neutralization potency *(7)*. **c.)** Competitive binding between IgG and ACE2 was performed by labeled yeast displaying aglycosylated S RBD with 10 μg/mL of indicated mAbs followed by labeling with biotinylated ACE2. **d.)** Single site saturation mutagenesis S RBD libraries were sorted by FACS using a competition experiment. The top cytogram shows the cell population collected for the control population without ACE2 labeling, while the bottom cytogram shows the cell population enriched in mutations able to bind ACE2 in the presence of a competing IgG. The specific cytogram shown is for nAb CC12.3 using the S RBD library corresponding to mutations at positions 437-537. **e.)** Per-mutation enrichment ratio distributions as a function of average depth of coverage control (top) and CC12.3 nAb competing experiment (bottom). **f.)** Heatmap showing predicted S RBD escape mutants for CC12.3 in blue. White cells are mutations with a p value for an FDR > 1, while grey cells are mutations not present in the mutational library. S RBD positions directly involved in binding ACE2 are colored gray. **g.)** Comparison of enrichment ratios (ER) for individual hits for CC12.3 vs. CC12.1. Closed circles represent escape mutant hits for both nAbs, whereas open circlers are escape mutant hits for only one nAb. **h.)** Solved structure of nAB CC12.3 in complex with S RBD (PDB ID 7KN6). **i.)** PSV neutralization curves for CC12.1, CC12.3 and CC6.29 on SARS-CoV-2 (top) and SARS-CoV-2 E484K (bottom). **j.)** PSV IC50 analysis for CC12.1, CC12.3 and CC6.29 on different identified mutations.

Next, we evaluated the ability of the mAb panel to competitively inhibit ACE2 binding to aglycosylated S RBD in an assay conceptually similar to the one previously described by Tan et al. *(13)*. Yeast displaying aglycosylated S RBD was first labeled with a saturating concentration of a given mAb and then co-incubated with biotinylated ACE2. Six mAbs completely ablated ACE2 binding, one mAb partially inhibited ACE2, and the remaining four did not prevent ACE2 binding (**Fig. 1c**). A direct correlation was observed between the previously determined neutralization potency of the antibody *(6, 8)* and the fluorescence signal increase in the competition assay (**Fig. 1c**). We conclude from these experiments that, excluding the S309 epitope, the aglycosylated S RBD platform faithfully recapitulates binding interactions of nAbs with S RBD *(7)*.

Our strategy for identifying potential S RBD escape mutants was as follows. First, we constructed a saturation mutagenesis library of aglycosylated S RBD containing all possible single missense and nonsense mutations for the 119 surface exposed positions of the RBD (96% coverage of the 2,380 possible library members; **Table S1** contains library coverage statistics) *(14)*. We labeled yeast displaying these RBD variants with a saturating concentration of nAb and then co-incubated with a saturating concentration of biotinylated ACE2. We then used fluorescence activated cell sorting (FACS) to screen for mutants that could bind ACE2, indicating that the RBD mutation allows for evasion of the nAb while not disrupting the ACE2 interaction critical for cell entry **(Fig. 1d, Fig. S3-S4**). Importantly, a control with no ACE2 labeling was sorted to set an empirical false discovery rate (FDR) for putative escape mutant hits (**Fig. 1d, Fig. S4**). Plasmid DNA from sorted cells were prepped and deep sequenced. We determined the enrichment ratio *(15)* for each mutant in the sorted population relative to a reference population, and then used the control population to set the FDR (**Fig. 1e, Fig S5**). We screened five different nAbs identified earlier as having completely ablated ACE2 binding (CC6.29, CC6.31, CC12.1, CC12.3, CC12.13). In all, we identified a total of 97 S RBD mutants that can escape recognition by at least one nAb (**Table S2**).

For all five nAbs, the putative escape mutant hits were localized in specific locations within the S RBD primary sequence (**Fig. 1f, Fig. S6-S10**). CC12.1 and CC12.3 belong to the public germline class VH3-53 *(6*, *8, 16)* and are representative of the subset of VH3-53 public antibodies with relatively short CDRH3 lengths *(17)*. Strikingly, these two nAbs share over 90% of the same RBD escape mutants (**Fig. 1g**), even though the light chain differs between the nAb. Structural complexes of antibodies CC12.1 and CC12.3 were previously solved in complex with S RBD *(18)*, affording a structural basis of individual escape mutants. Escape mutants for both of the VH3-53 nAbs CC12.1 and CC12.3 clustered at the same location on the S RBD mainly peripheral to the ACE2 binding site (**Fig 1h; Fig. S11**).

Having identified a number of putative escape mutants from the mutagenesis library screening we sought to determine how this functional screening correlated with the more conventional pseudovirus neutralization assay. A panel of MLV-based SARS-CoV-2 pseudoviruses were generated that contained single mutations predicted by the mutagenesis scanning to allow escape from one of the antibodies screened, as well as several irrelevant control mutations. Antibodies CC12.1, CC12.3 and CC6.29 were screened against the original SARS-CoV-2 pseudovirus as well as this panel of mutant pseudoviruses in duplicate (**Fig. 1i**), and the resulting IC_50_s were compared to calculate the effect on antibody neutralization potency (**Fig. 1j; Fig. S12**). Consistent with RBD mutagenesis library and structural analysis, CC6.29 failed to neutralize F486I, E484K, and T478R variants. Additionally, K417N, K417T, and D420K hotspot mutants completely escaped neutralization for both CC12.1 and CC12.3. The only instance we tested where the mutagenesis scanning data differed from the pseudovirus results was at N501Y that was predicted to confer escape from CC12.1 and CC12.3 but had no effect on the *in vitro* neutralization potency. Although it is unclear why this discrepancy occurred, we note that N501Y significantly increases the affinity of RBD for ACE2, which could result in ACE2 competing off bound nAb.

Finally, we performed biological replicates where the mutagenesis library corresponding to S RBD positions 437-537 was separately transformed into yeast and screened against nAbs CC6.29, CC12.1, and CC12.3. While the overall magnitude of the enrichment ratios were lower than in the initial experiment, nearly the same set of escape mutants were identified for CC6.29, and escape mutants originally identified for all nAbs had significantly higher ERs than other variants in the replicate (p-value range 4.2e-4 to 1.9e-11, one-sided Welch’s t-test) (**Fig. S13**).

Selected per-position heatmaps, and structural mapping of S RBD escape mutants, are shown in **Fig. 2** for all five nAbs. Closer examination of these datasets reveals key features of the RBD escape mutant response. CC12.1 and CC12.3 nAbs share over 90% of the same RBD escape mutants (**Fig. 1g**), including notable hotspot mutations occurring at K417, D420, Y421, and Q498 (**Fig. 2a**). Interestingly, multiple aromatic substitutions at Q498 escape recognition for CC12.1 & CC12.3 even though the antibodies have different light chains and recognition motifs for that position. Introduction of an aromatic residue at Q498 introduces substantial van der Waals clashes that are likely unresolved without antibody loop movement. The other VH3-53 nAb tested, CC12.13, has a 15 amino acid length CDRH3 that likely has a distinct binding mode than that for CC12.1 and CC12.3 *(17)*. Consistent with this, the CC12.13 escape mutants identified are mostly different from those for CC12.1/CC12.3 (**Fig. S10**).

**Figure 2.**
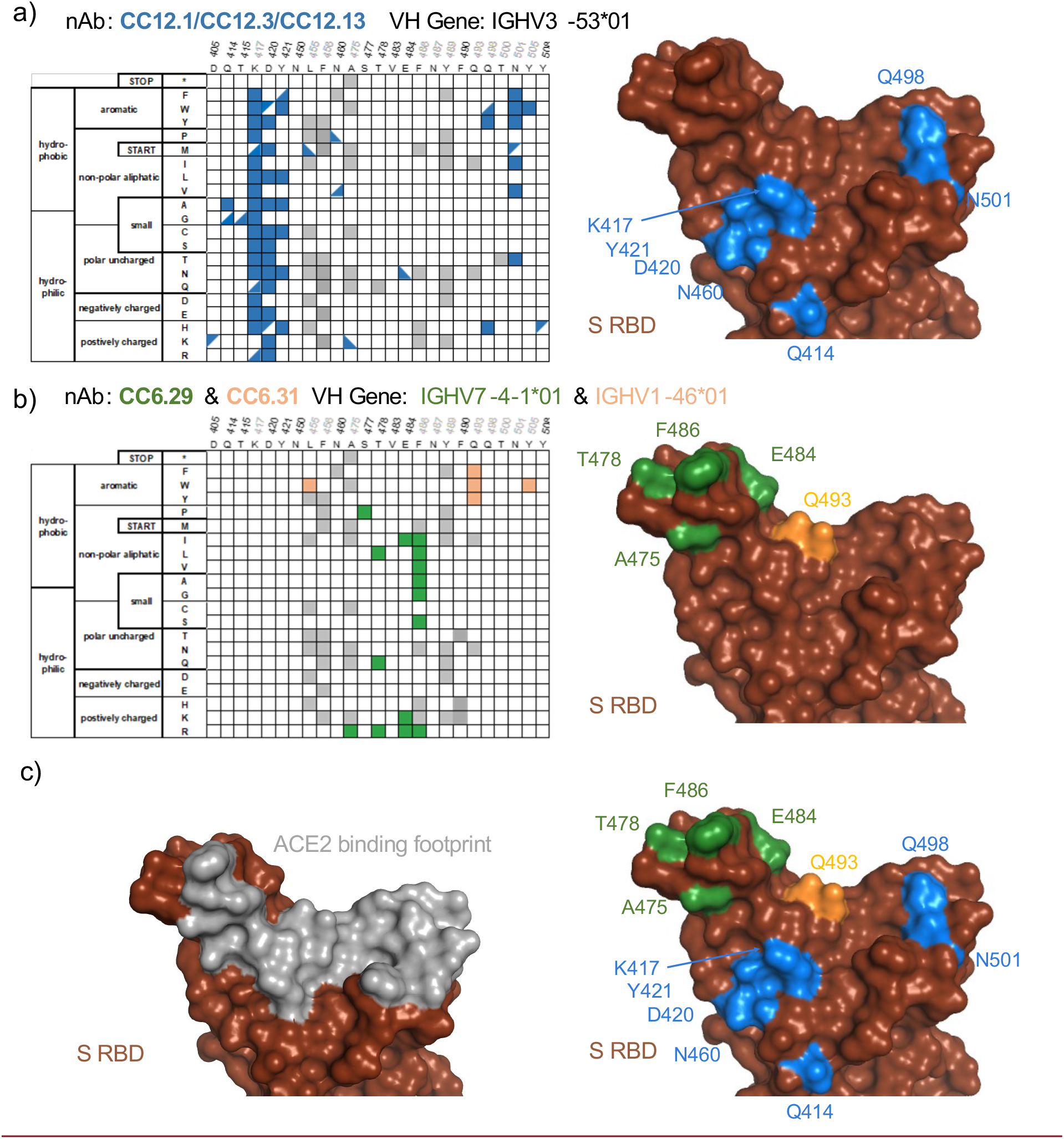
Sequence determinants and structural basis of S RBD escape mutants. **a-c)** (left) Limited per-position heatmap and (right) mutations mapped onto the S RBD-ACE2 structural complex (PDB ID: 6M0J). For clarity, only positions with two or more escape mutations are shown with surface colored. Panels are for **a)** nAbs CC12.1, CC12.3, and CC12.13. Boxes indicate escape mutants for two or more nAbs, while triangles indicate an escape mutant identified for just one nAb (top left: CC12.1, bottom right: CC12.3, bottom left: CC12.13); **b)** CC6.29 and CC6.31 (orange); **c)** overlay of escape mutants from all nAbs onto the S RBD-ACE2 structural complex.

Another nAb screened, CC6.29, has a completely different escape mutant profile compared with CC12.1/CC12.3. The 15 potential RBD escape mutants for CC6.29 center around the structural ‘knob’ of positions A475, S477, T478, E484, and F486 (**Fig. 2b**). E484K shared by the B.1.351 and B.1.526 lineages is identified as an escape mutant for this nAb, but the structurally adjacent S477N mutation newly identified in the B.1.526 lineage does not escape CC6.29 neutralization. Intriguingly, S477P is identified as an escape mutant for this nAb. F486 shows up as a mutational hotspot even though that position is involved in the recognition of ACE2. However, a previous mutational scan of S RBD shows that F486 mutation does not significantly impact ACE2 binding affinity *(10)*. CC6.31 escape mutants partially overlap with CC6.29 but implicate a different set of mutants (**Fig. 2b**). Multiple mutations at Q493 escape CC6.31, including Q493 substitutions to aromatic positions F/W.

In total, the five nAbs map a partially overlapping surface with the ACE2 binding site that is primed for antibody escape. In comparison with the binding footprint of ACE2 **(Fig. 2c)**, the escape mutants almost completely map to the outer binding shell and periphery of the interaction surface, akin to an O ring circumnavigating the receptor binding site. Out of the identified escape mutants, residues K417, F486, Q493, N501, and Y505 are located on the ACE2 footprint (**Fig. 2c**). While mutations on K417 and F486 do not significantly change the RBD affinity to ACE2, mutations on N501 can both increase or decrease affinity depending on the substitution. The Y505W mutant shared by CC6.31, CC12.1, and CC12.3 also increases ACE2 affinity *(10)*.

We puzzled why mutations at D420 were so deleterious to the neutralization potency of the VH3-53 nAbs given that this residue is on the outer periphery of the binding epitope. To obtain insight into this question, we performed 100 ns aqueous molecular dynamics (MD) simulations of CC12.1 and CC12.3 in complex with wildtype S RBD and S RBD incorporated with the D420E, D420K, or the Y421N mutation (see SI for details). In the control simulation with CC12.1, D420 on the RBD and CDRH2 S56 on CC12.1 form persistent hydrogen bonds, and Y421 on the RBD is tightly bound within a pocket of CC12.1 residues (**Fig. 3a**). With the D420E mutation, the increased length of E420 disrupts its ability to hydrogen bond with S56, requiring it to adopt a bent conformation (**Fig. 3b**). This forces Y421 out of the antibody pocket, causing increased fluctuations in neighboring RBD loops which persist throughout the entire 100 ns production simulation (**Fig. S14a, b**). With the D420K mutation, hydrogen bonding with S56 is completely disrupted, and with the Y421N mutation, N421 is too short to interact with the antibody pocket (**Fig. S14c**). Similar escape mechanisms are observed for CC12.3 with all three RBD mutations, including increased fluctuations at one of the same key sites (K458) on the RBD in response to the D420E mutation (**Fig. S14d, e**).

**Figure 3.**
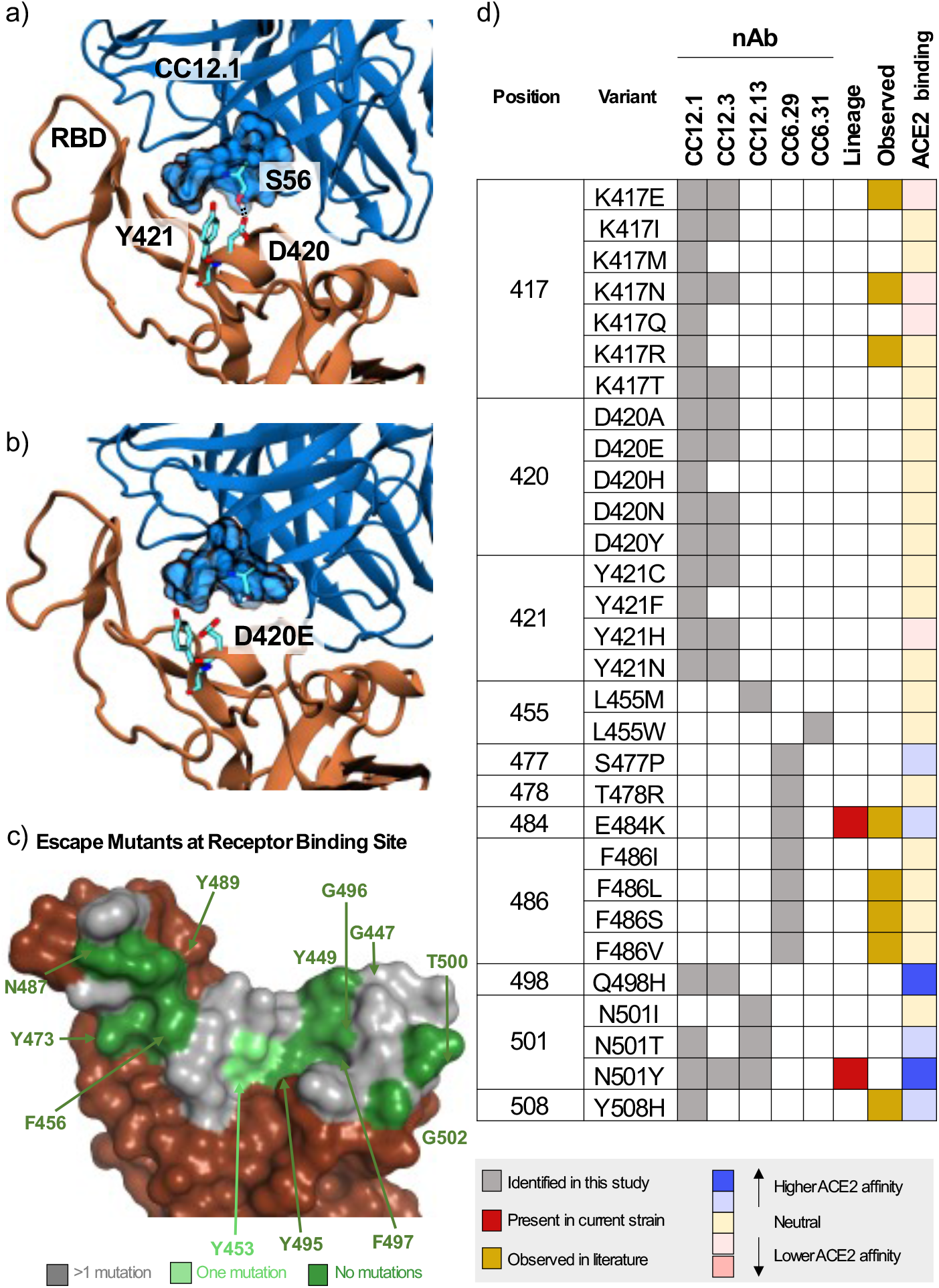
Mechanistic, structural, and sequence analysis of SARS-CoV-2 escape mutants. **a-b)** Snapshots from MD trajectories showing **a.)** key interactions in the control simulation of S RBD in complex with CC12.1, and **b.)** mechanism of escape of S RBD from CC12.1 due to the D420E mutation. Images were rendered with Visual Molecular Dynamics (VMD *(25)*), and black dotted lines indicate persistent hydrogen bonds. **c.)** S RBS positions are colored by the number of escape mutants identified to date. RBS residues involving the S RBD-ACE2 structural complex (PDB ID: 6M0J) are colored by number of escape mutants identified to date. **d.)** Summary of 1-nt escape mutants identified in the present study. Lineage column indicates presence of the given mutation amongst currently circulating SARS-CoV-2 strains, while the observed column refers to an escape mutant previously identified in literature *(11, 19–22)*. ACE2 binding indicates affinity to ACE2 based on the measurements by Starr et al. *(10)*.

There have been a number of recent approaches to identify specific S escape mutants (summarized in **Table S3**) *(11, 19–22)*. A survey of the existing escape mutant literature, along with escape mutants identified in the present work, allows us to identify the absolute and near-absolute escape resistant ACE2 receptor binding site (RBS) residues in the context of the original lineage (**Fig. 3c**). One resistant patch is found around F456/Y473/N487/Y489 while other residues are discontinuous patches on the remainder of the RBS. We note that many of these same resistant residues are identical to those from SARS-CoV (Y449, N487, Y489, G496, T500 and G502). The lack of a contiguous surface at the RBS that is conserved makes it highly unlikely that one could identify a naïve nAb targeting the RBS that is completely resistant to escape.

A major near-term concern with public health implications are identification of the set of single nucleotide polymorphisms that encode for escape mutants on the S RBD. A summary of 1-nucleotide (nt) escape mutants identified in the present work is shown in **Fig. 3d**. To our knowledge, 22/35 (63%) of 1-nt escape mutants identified from this nAb panel have not previously been identified, including hotspot positions D420 and Y421 that escape recognition by the abundant VH3-53 nAbs. Other notable residues identified here include S477, Q498, and Y501, as these positions lie directly on the receptor binding site and all have been shown to slightly increase binding affinity to ACE2 *(10)*. Mutants E484K and N501Y in currently circulating lineages escape some but not all of the nAbs on the panel.

We have developed a yeast platform that allows for the rapid identification of SARS-CoV-2 S RBD escape mutants for a given nAb. While other platforms to identify escape mutants have recently been described, key advantages of the approach presented here includes: (i.) screening by competitive binding against ACE2 which more precisely mimics how actual viral infection can still persist despite antibody binding; (ii.) a robust and rigorous hit identification algorithm; (iii.) a safe working environment, as it does not use live virus; and (iv.) a relatively fast identification, as the RBD library can be screened against a given nAb and analyzed in under a week. There also exist drawbacks. First, the present method is limited to mapping escape mutants for anti-S-RBD nAbs that directly compete with ACE2 for binding. Many nAbs neutralize by targeting S epitopes across protomers *(23)* or on the N-terminal domain *(24)*, and a robust platform for S ectodomain display would enable more comprehensive studies. We attempted to develop a yeast platform for S ectodomain but were unsuccessful: we screened media composition, expression temperature, protein orientation (**Fig. S15**), and mutations (1,909 mutants screened with only two potential hits) (**Data S2-3, Table S1, Fig. S16**). Second, the presented assay measures the ability of a given mutant to escape nAb blockade of ACE2. While from all available data the assay appears to correlate well in the context of pseudo-virus, each mutation is pleiotropic with unknown fitness effects beyond escape for a given nAb; the true RBS escape mutants that do not appreciably impede viral fitness will be a subset of the mutations identified here.

Still, using this method we were able to identify specific failure mechanisms for five different nAbs. This tool can be easily adapted and contribute to developing the next generation of broadly neutralizing antibodies against SARS-CoV-2, as well as suggest mutations to include for the next generation of vaccines. To that end, it would be interesting to see whether our yeast platform presented here is robust enough to identify escape mutants from bulk sera from convalescent or vaccinated individuals.

## Acknowledgements

We thank scientists K. Jackson and A. Scott at the BioFrontiers sequencing core for technical guidance, and members of the Whitehead lab for intellectual and material support, including Matt Bedewitz for reagent prep. The anti-SARS-CoV-2 RBD antibody panel used was a kind gift from Dennis Burton’s lab at Scripps, and the ACE2-Fc and CR3022 were kind gifts from Neil King’s lab at the University of Washington.

## Author Contributions

Designed experiments: IFU, PJS, MBK, ERR, KGS, JJ, TAW; Performed experiments: IFU, PJS, MBK, SB, LP, FZ, ERR; Performed simulations: ERR, KGS; Developed algorithms and software: PJS, CMH, TAW; Wrote paper: IFU, PJS, MBK, KGS, TAW.

## Data and Reagent Availability

Raw sequencing reads for this work have been deposited in the SRA (Accession #s SAMN18250431-SAMN18250483). All plasmids and mutational libraries used in this work are available from AddGene (AddGene Collection **To Be Added Upon Publication**). All scripts used to process and analyze deep sequencing data are freely available on Github (https://github.com/WhiteheadGroup/SpikeRBDStabilization.git).

