## Supplemental Information for "One-shot identification of SARS-CoV-2 S RBD escape mutants using yeast screening"

3415 Colorado Avenue, Boulder, CO 80305

**One Sentence Summary:** We present a facile method to identify antibody escape mutants on SARS-CoV-2 S RBD.

#### ABSTRACT

The potential emergence of SARS-CoV-2 Spike (S) escape mutants is a threat to reduce the efficacy of existing vaccines and neutralizing antibody (nAb) therapies. An understanding of the antibody/S escape mutations landscape is urgently needed to preemptively address this threat. Here we describe a rapid method to identify escape mutants for nAbs targeting the S receptor binding site. We identified escape mutants for five nAbs, including three from the public germline class VH3-53 elicited by natural COVID-19 infection. Escape mutations predominantly mapped to the periphery of the ACE2 recognition site on the RBD with K417, D420, Y421, F486, and Q493 as notable hotspots. We provide libraries, methods, and software as an openly available community resource to accelerate new therapeutic strategies against SARS-CoV-2.

### MAIN

The type I viral fusion protein Spike (S) is a major antigenic determinant of SARS-CoV-2 and is the antigen used in all approved COVID-19 vaccines *(1–3)*. Recently, the B.1.1.7 (N501Y; U.K.), B.1.351 (E484K; South Africa), B.1.427 (L452R; California), and B.1.526 (S477N, E484K; NY) viral lineages have emerged. All encode single nucleotide substitutions in the S receptor binding domain (RBD) near recognition site for its cellular target angiotensin-converting enzyme 2 (ACE2) *(3–5)*.

Dozens of studies have reported the structural, epitopic, and functional landscape of non-neutralizing monoclonal antibodies and nAbs targeting trimeric S *(6–8)*. A prophetic understanding of the mutations on S that could evade antibody recognition would enable development of better vaccine boosters and monoclonal antibody therapies. Thus, we sought to develop an S RBD yeast surface display (YSD) platform (**Fig. S1**) *(9)*, as we hypothesized that broad identification of SARS-CoV-2 S escape mutants could be found by integrating high throughput screening platforms with deep sequencing. While a similar platform uses the loss of nAb binding to identify escape mutants *(10, 11)*, we rationalized that a functional screening assay that directly measures the ability of a nAb to compete with ACE2 for S RBD binding, would be a comparatively strong predictor of RBD escapability, as it accounts for mutations in RBD that would disrupt S binding to ACE2.

We had previously developed an aglycosylated S-RBD YSD platform (S RBD(333-537)-N343Q) *(8)* that can bind specifically to ACE2 (**Fig. 1a**). This S RBD construct has its one native N-linked glycan removed (N343Q) as the heavy N-linked mannosylation endemic of *S. cerevisiae* could hamper anti-S RBD mAb recognition. Cell surface titrations of CR3022 IgG and nAb HKU-910-30 IgG yielded apparent dissociation constants comparable to reported *in vitro* results *(6,8)* (**Fig. S2**). We next tested a panel of eleven additional anti-S RBD mAbs for binding to aglycosylated RBD *(7)*. Ten of the eleven mAbs recognized aglycosylated S RBD (**Fig. 1b**). The one panel member that did not bind, CC6.33, selectively recognizes the S309 epitope on the RBD containing the N-linked glycan at position 343 *(12)*.

Next, we evaluated the ability of the mAb panel to competitively inhibit ACE2 binding to aglycosylated S RBD in an assay conceptually similar to the one previously described by Tan et al. *(13)*. Yeast displaying aglycosylated S RBD was first labeled with a saturating concentration of a given mAb and then co-incubated with biotinylated ACE2. Six mAbs completely ablated ACE2 binding, one mAb partially inhibited ACE2, and the remaining four did not prevent ACE2 binding (**Fig. 1c**). A direct correlation was observed between the previously determined neutralization potency of the antibody *(6, 8)* and the fluorescence signal increase in the competition assay (**Fig. 1c**). We conclude from these experiments that, excluding the S309 epitope, the aglycosylated S RBD platform faithfully recapitulates binding interactions of nAbs with S RBD *(7)*.

Our strategy for identifying potential S RBD escape mutants was as follows. First, we constructed a saturation mutagenesis library of aglycosylated S RBD containing all possible single missense and nonsense mutations for the 119 surface exposed positions of the RBD (96% coverage of the 2,380 possible library members; **Table S1** contains library coverage statistics) *(14)*. We labeled yeast displaying these RBD variants with a saturating concentration of nAb and then co-incubated with a saturating concentration of biotinylated ACE2. We then used fluorescence activated cell sorting (FACS) to screen for mutants that could bind ACE2, indicating that the RBD mutation allows for evasion of the nAb while not disrupting the ACE2 interaction critical for cell entry **(Fig. 1d, Fig. S3-S4)**. Importantly, a control with no ACE2 labeling was sorted to set an empirical false discovery rate (FDR) for putative escape mutant hits (**Fig. 1d, Fig. S4**). Plasmid DNA from sorted cells were prepped and deep sequenced. We determined the enrichment ratio *(15)* for each mutant in the sorted population relative to a reference population, and then used the control population to set the FDR (**Fig. 1e, Fig S5**). We screened five different nAbs identified earlier as having completely ablated ACE2 binding (CC6.29, CC6.31, CC12.1, CC12.3, CC12.13). In all, we identified a total of 97 S RBD mutants that can escape recognition by at least one nAb (**Table S2**).

For all five nAbs, the putative escape mutant hits were localized in specific locations within the S RBD primary sequence (**Fig. 1f, Fig. S6-S10**). CC12.1 and CC12.3 belong to the public germline class VH3-53 *(6 ,8, 16)* and are representative of the subset of VH3-53 public antibodies with relatively short CDRH3 lengths *(17)*. Strikingly, these two nAbs share over 90% of the same RBD escape mutants (**Fig. 1g**), even though the light chain differs between the nAb. Structural complexes of antibodies CC12.1 and CC12.3 were previously solved in complex with S RBD *(18)*, affording a structural basis of individual escape mutants. Escape mutants for both of the VH3-53 nAbs CC12.1 and CC12.3 clustered at the same location on the S RBD mainly peripheral to the ACE2 binding site (**Fig 1h; Fig. S11**).

Having identified a number of putative escape mutants from the mutagenesis library screening we sought to determine how this functional screening correlated with the more conventional pseudovirus neutralization assay. A panel of MLV-based SARS-CoV-2 pseudoviruses were generated that contained single mutations predicted by the mutagenesis scanning to allow escape from one of the antibodies screened, as well as several irrelevant control mutations. Antibodies CC12.1, CC12.3 and CC6.29 were screened against the original SARS-CoV-2 pseudovirus as well as this panel of mutant pseudoviruses in duplicate (**Fig. 1i**), and the resulting $IC_{50}$s were compared to calculate the effect on antibody neutralization potency (**Fig. 1j; Fig. S12**). Consistent with RBD mutagenesis library and structural analysis, CC6.29 failed to neutralize F486I, E484K, and T478R variants. Additionally, K417N, K417T, and D420K hotspot mutants completely escaped neutralization for both CC12.1 and CC12.3. The only instance we tested where the mutagenesis scanning data differed from the pseudovirus results was at N501Y that was predicted to confer escape from CC12.1 and CC12.3 but had no effect on the *in vitro* neutralization potency. Although it is unclear why this discrepancy occurred, we note that N501Y significantly increases the affinity of RBD for ACE2, which could result in ACE2 competing off bound nAb.

Finally, we performed biological replicates where the mutagenesis library corresponding to S RBD positions 437-537 was separately transformed into yeast and screened against nAbs CC6.29, CC12.1, and CC12.3. While the overall magnitude of the enrichment ratios were lower than in the initial experiment, nearly the same set of escape mutants were identified for CC6.29, and escape mutants originally identified for all nAbs had significantly higher ERs than other variants in the replicate (p-value range 4.2e-4 to 1.9e-11, one-sided Welch's t-test) (**Fig. S13**).

Selected per-position heatmaps, and structural mapping of S RBD escape mutants, are shown in **Fig. 2** for all five nAbs. Closer examination of these datasets reveals key features of the RBD escape mutant response. CC12.1 and CC12.3 nAbs share over 90% of the same RBD escape mutants (**Fig. 1g**), including notable hotspot mutations occurring at K417, D420, Y421, and Q498 (**Fig. 2a**). Interestingly, multiple aromatic substitutions at Q498 escape recognition for CC12.1 & CC12.3 even though the antibodies have different light chains and recognition motifs for that position. Introduction of an aromatic residue at Q498 introduces substantial van der Waals clashes that are likely unresolved without antibody loop movement. The other VH3-53 nAb tested, CC12.13, has a 15 amino acid length CDRH3 that likely has a distinct binding mode than that for CC12.1 and CC12.3 *(17)*. Consistent with this, the CC12.13 escape mutants identified are mostly different from those for CC12.1/CC12.3 (**Fig. S10**).

Another nAb screened, CC6.29, has a completely different escape mutant profile compared with CC12.1/CC12.3. The 15 potential RBD escape mutants for CC6.29 center around the structural 'knob' of positions A475, S477, T478, E484, and F486 (**Fig. 2b**). E484K shared by the B.1.351 and B.1.526 lineages is identified as an escape mutant for this nAb, but the structurally adjacent S477N mutation newly identified in the B.1.526 lineage does not escape CC6.29 neutralization. Intriguingly, S477P is identified as an escape mutant for this nAb. F486 shows up as a mutational hotspot even though that position is involved in the recognition of ACE2. However, a previous mutational scan of S RBD shows that F486 mutation does not significantly impact ACE2 binding affinity *(10)*. CC6.31 escape mutants partially overlap with CC6.29 but implicate a different set of mutants (**Fig. 2b**). Multiple mutations at Q493 escape CC6.31, including Q493 substitutions to aromatic positions F/W.

In total, the five nAbs map a partially overlapping surface with the ACE2 binding site that is primed for antibody escape. In comparison with the binding footprint of ACE2 **(Fig. 2c)**, the escape mutants almost completely map to the outer binding shell and periphery of the interaction surface, akin to an O ring circumnavigating the receptor binding site. Out of the identified escape mutants, residues K417, F486, Q493, N501, and Y505 are located on the ACE2 footprint (**Fig. 2c**). While mutations on K417 and F486 do not significantly change the RBD affinity to ACE2, mutations on N501 can both increase or decrease affinity depending on the substitution. The Y505W mutant shared by CC6.31, CC12.1, and CC12.3 also increases ACE2 affinity *(10)*.

We puzzled why mutations at D420 were so deleterious to the neutralization potency of the VH3-53 nAbs given that this residue is on the outer periphery of the binding epitope. To obtain insight into this question, we performed 100 ns aqueous molecular dynamics (MD) simulations of CC12.1 and CC12.3 in complex with wildtype S RBD and S RBD incorporated with the D420E, D420K, or the Y421N mutation (see SI for details). In the control simulation with CC12.1, D420 on the RBD and CDRH2 S56 on CC12.1 form persistent hydrogen bonds, and Y421 on the RBD is tightly bound within a pocket of CC12.1 residues (**Fig. 3a**). With the D420E mutation, the increased length of E420 disrupts its ability to hydrogen bond with S56, requiring it to adopt a bent conformation (**Fig. 3b**). This forces Y421 out of the antibody pocket, causing increased fluctuations in neighboring RBD loops which persist throughout the entire 100 ns production simulation (**Fig. S14a, b**). With the D420K mutation, hydrogen bonding with S56 is completely disrupted, and with the Y421N mutation, N421 is too short to interact with the antibody pocket (**Fig. S14c**). Similar escape mechanisms are observed for CC12.3 with all three RBD mutations, including increased fluctuations at one of the same key sites (K458) on the RBD in response to the D420E mutation (**Fig. S14d, e**).

There have been a number of recent approaches to identify specific S escape mutants (summarized in **Table S3**) *(11, 19–22)*. A survey of the existing escape mutant literature, along with escape mutants identified in the present work, allows us to identify the absolute and near-absolute escape resistant ACE2 receptor binding site (RBS) residues in the context of the original lineage (**Fig. 3c**). One resistant patch is found around F456/Y473/N487/Y489 while other residues are discontinuous patches on the remainder of the RBS. We note that many of these same resistant residues are identical to those from SARS-CoV (Y449, N487, Y489, G496, T500 and G502). The lack of a contiguous surface at the RBS that is conserved makes it highly unlikely that one could identify a naïve nAb targeting the RBS that is completely resistant to escape.

A major near-term concern with public health implications are identification of the set of single nucleotide polymorphisms that encode for escape mutants on the S RBD. A summary of 1-nucleotide (nt) escape mutants identified in the present work is shown in **Fig. 3d**. To our knowledge, 22/35 (63%) of 1-nt escape mutants identified from this nAb panel have not previously been identified, including hotspot positions D420 and Y421 that escape recognition by the abundant VH3-53 nAbs. Other notable residues identified here include S477, Q498, and Y501, as these positions lie directly on the receptor binding site and all have been shown to slightly increase binding affinity to ACE2 *(10)*. Mutants E484K and N501Y in currently circulating lineages escape some but not all of the nAbs on the panel.

We have developed a yeast platform that allows for the rapid identification of SARS-CoV-2 S RBD escape mutants for a given nAb. While other platforms to identify escape mutants have recently been described, key advantages of the approach presented here includes: (i.) screening by competitive binding against ACE2 which more precisely mimics how actual viral infection can still persist despite antibody binding; (ii.) a robust and rigorous hit identification algorithm; (iii.) a safe working environment, as it does not use live virus; and (iv.) a relatively fast identification, as the RBD library can be screened against a given nAb and analyzed in under a week. There also exist drawbacks. First, the present method is limited to mapping escape mutants for anti-S-RBD nAbs that directly compete with ACE2 for binding. Many nAbs neutralize by targeting S epitopes across protomers *(23)* or on the N-terminal domain *(24)*, and a robust platform for S ectodomain display would enable more comprehensive studies. We attempted to develop a yeast platform for S ectodomain but were unsuccessful: we screened media composition, expression temperature, protein orientation (**Fig. S15**), and mutations (1,909 mutants screened with only two potential hits) (**Data S2-3, Table S1, Fig. S16**). Second, the presented assay measures the ability of a given mutant to escape nAb blockade of ACE2. While from all available data the assay appears to correlate well in the context of pseudo-virus, each mutation is pleiotropic with unknown fitness effects beyond escape for a given nAb; the true RBS escape mutants that do not appreciably impede viral fitness will be a subset of the mutations identified here.

Still, using this method we were able to identify specific failure mechanisms for five different nAbs. This tool can be easily adapted and contribute to developing the next generation of broadly neutralizing antibodies against SARS-CoV-2, as well as suggest mutations to include for the next generation of vaccines. To that end, it would be interesting to see whether our yeast platform presented here is robust enough to identify escape mutants from bulk sera from convalescent or vaccinated individuals.

#### Acknowledgements

We thank scientists K. Jackson and A. Scott at the BioFrontiers sequencing core for technical guidance, and members of the Whitehead lab for intellectual and material support, including Matt Bedewitz for reagent prep. The anti-SARS-CoV-2 RBD antibody panel used was a kind gift from Dennis Burton's lab at Scripps, and the ACE2-Fc and CR3022 were kind gifts from Neil King's lab at the University of Washington. **Funding:** Research reported in this publication was supported by the National Institute Of Allergy And Infectious Diseases of the National Institutes of Health under Award Number R01AI141452 to T.A.W. The content is solely the responsibility of the authors and does not necessarily represent the official views of the National Institutes of Health. This research was also supported by a Balsells Fellowship to I.F.U. and a CU Boulder BSI fellowship to C.M.H. **Author Contributions:** Designed experiments: IFU, PJS, MBK, ERR, KGS, JJ, TAW; Performed experiments: IFU, PJS, MBK, SB, LP, FZ, ERR; Performed simulations: ERR, KGS; Developed algorithms and software: PJS, CMH, TAW; Wrote paper: IFU, PJS, MBK, KGS, TAW. **Competing Interests:** IFU, PJS, MBK, CH, JJ, TAW declare competing interests.

#### Data and Reagent Availability

Raw sequencing reads for this work have been deposited in the SRA (Accession #s SAMN18250431-SAMN18250483). All plasmids and mutational libraries used in this work are available from AddGene (AddGene Collection **To Be Added Upon Publication**). All scripts used to process and analyze deep sequencing data are freely available on Github (https://github.com/WhiteheadGroup/SpikeRBDStabilization.git).

### Figures

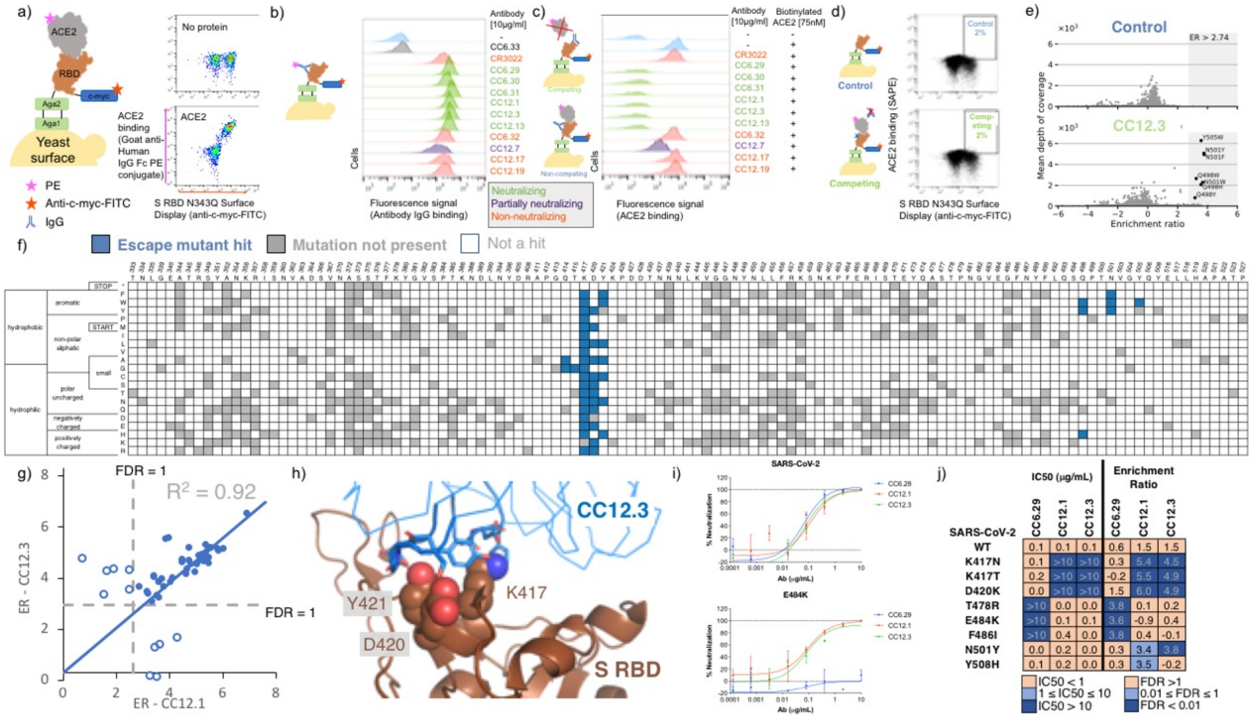


**Figure 1.** Identification and validation of SARS-CoV-2 S RBD escape mutants using yeast screening. **a)** Cartoon of the yeast display construct S-RBD(333-537)-N343Q. Cytograms show specific binding in the presence, but not absence, of ACE2-Fc. **b)** Binding profiles of aglycosylated S RBD labeled with 10 $\mu$g/mL of indicated mAb. Antibodies are color coded according to neutralization potency *(7)*. **c.)** Competitive binding between IgG and ACE2 was performed by labeled yeast displaying aglycosylated S RBD with 10 $\mu$g/mL of indicated mAbs followed by labeling with biotinylated ACE2. **d.)** Single site saturation mutagenesis S RBD libraries were sorted by FACS using a competition experiment. The top cytogram shows the cell population collected for the control population without ACE2 labeling, while the bottom cytogram shows the cell population enriched in mutations able to bind ACE2 in the presence of a competing IgG. The specific cytogram shown is for nAb CC12.3 using the S RBD library corresponding to mutations at positions 437-537. **e.)** Per-mutation enrichment ratio distributions as a function of average depth of coverage control (top) and CC12.3 nAb competing experiment (bottom). **f.)** Heatmap showing predicted S RBD escape mutants for CC12.3 in blue. White cells are mutations with a p value for an FDR > 1, while grey cells are mutations not present in the mutational library. S RBD positions directly involved in binding ACE2 are colored gray. **g.)** Comparison of enrichment ratios (ER) for individual hits for CC12.3 vs. CC12.1. Closed circles represent escape mutant hits for both nAbs, whereas open circlers are escape mutant hits for only one nAb. **h.)** Solved structure of nAB CC12.3 in complex with S RBD (PDB ID 7KN6). **i.)** PSV neutralization curves for CC12.1, CC12.3 and CC6.29 on SARS-CoV-2 (top) and SARS-CoV-2 E484K (bottom). **j.)** PSV IC50 analysis for CC12.1, CC12.3 and CC6.29 on different identified mutations.

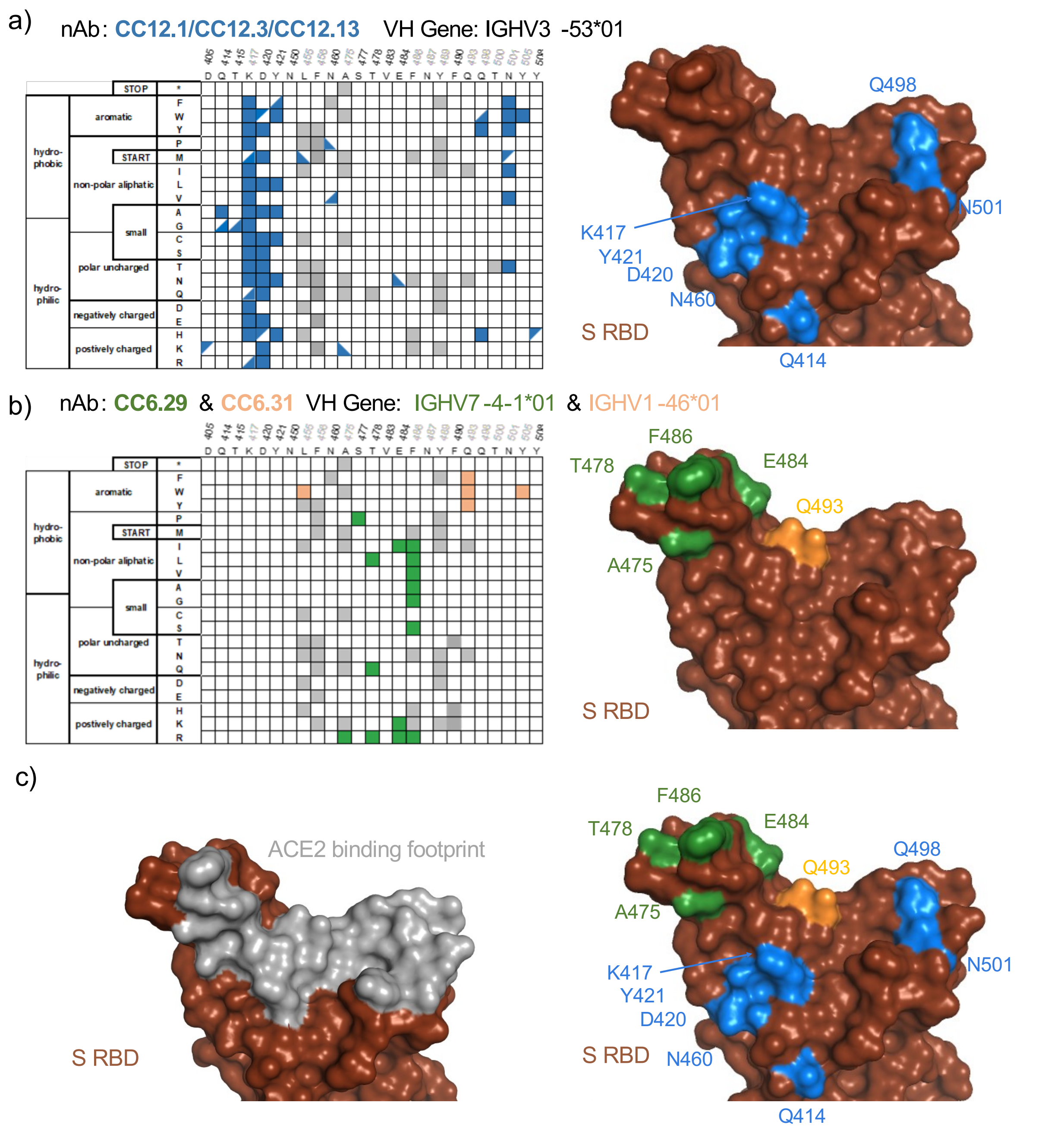


**Figure 2. Sequence determinants and structural basis of S RBD escape mutants. a-c)** (left) Limited per-position heatmap and (right) mutations mapped onto the S RBD-ACE2 structural complex (PDB ID: 6M0J). For clarity, only positions with two or more escape mutations are shown with surface colored. Panels are for **a)** nAbs CC12.1, CC12.3, and CC12.13. Boxes indicate escape mutants for two or more nAbs, while triangles indicate an escape mutant identified for just one nAb (top left: CC12.1, bottom right: CC12.3, bottom left: CC12.13) ; **b)** CC6.29 and CC6.31 (orange); **c)** overlay of escape mutants from all nAbs onto the S RBD-ACE2 structural complex.

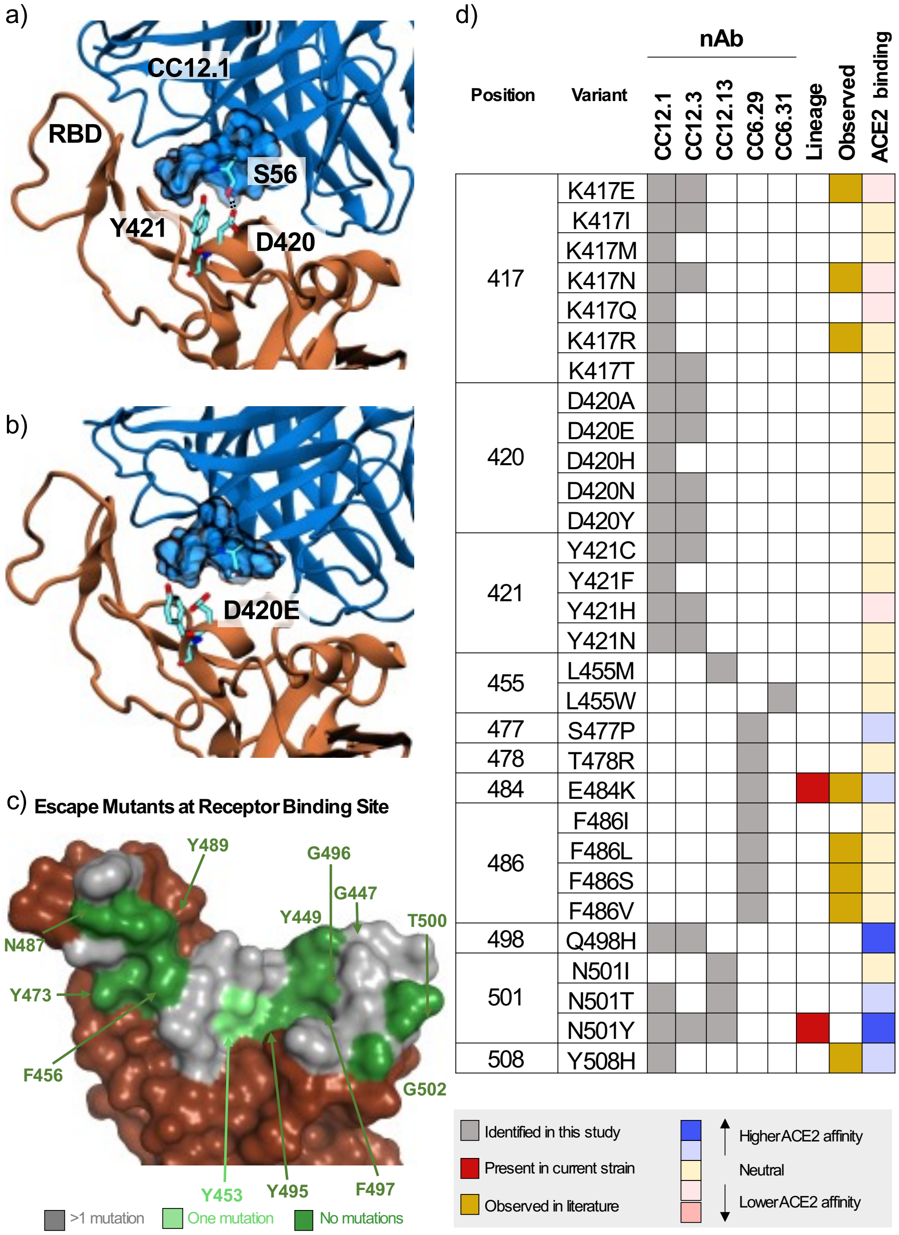


**Figure 3.** Mechanistic, structural, and sequence analysis of SARS-CoV-2 escape mutants. **a-b)** Snapshots from MD trajectories showing **a.)** key interactions in the control simulation of S RBD in complex with CC12.1, and **b.)** mechanism of escape of S RBD from CC12.1 due to the D420E mutation. Images were rendered with Visual Molecular Dynamics (VMD *(25)*), and black dotted lines indicate persistent hydrogen bonds. **c.)** S RBS positions are colored by the number of escape mutants identified to date. RBS residues involving the S RBD-ACE2 structural complex (PDB ID: 6M0J) are colored by number of escape mutants identified to date. **d.)** Summary of 1-nt escape mutants identified in the present study. Lineage column indicates presence of the given mutation amongst currently circulating SARS-CoV-2 strains, while the observed column refers to an escape mutant previously identified in literature *(11, 19–22)*. ACE2 binding indicates affinity to ACE2 based on the measurements by Starr et al. *(10)*.

Supplementary Materials for

### One-shot identification of SARS-CoV-2 S RBD escape mutants using yeast screening

Irene Francino Urdaniz[1,#], Paul J. Steiner[1,#], Monica B. Kirby[1,#], Fangzhu Zhao[2], Cyrus M. Haas[1], Shawn Barman[2], Emily R. Rhodes[1], Linghang Peng[2], Kayla G. Sprenger[1], Joseph G. Jardine[2,3], Timothy A. Whitehead[1,*]

[1]Department of Chemical and Biological Engineering, University of Colorado, Boulder, CO 80305, USA
[2]The Scripps Research Institute, La Jolla, CA, USA
[3]International AIDS Vaccine Initiative, New York, NY, USA

### Co-authors contributed equally.

**This PDF file includes:**

Materials and Methods
Figures S1 to S17
Tables S1 to S5
Captions for Data S1 to S3

**Other Supplementary Materials for this manuscript include the following:**

Data S1 to S3

#### Materials and Methods

##### Plasmid constructs

All plasmids used for this work are listed in **Table S4** and all primers in **Table S5**. All plasmids were verified by Sanger sequencing. Yeast display constructs for SARS-CoV-2 spike protein ectodomain (GenBank MN908947 with a GSAS substitution at the furin cleavage site (682-685) and proline substitutions at positions 986 and 987 (1), and a C-terminal T4 fibritin trimerization domain), as shown in **Figure S1**, were constructed as follows. Spike was codon optimized for *Saccharomyces cerevisiae* with Benchling software using default options, split into three gene blocks (hereafter labeled A, B, and C) each encoded with BsaI restriction sites with overhangs (2), synthesized as gBlocks (IDT), and cloned into pUC19 (Addgene: #50005) using SalI/KpnI restriction sites. This yielded the spike fragment entry plasmids pUC19-S-ecto-B and pUC19-S-ecto-C-Nterm. To construct pUC19-S-ecto-A-Nterm-KanR (the spike fragment destination plasmid), PCR was used to amplify both the kanamycin resistance gene from pETconNK (Addgene: #81169) with primers MBK-175 and MBK-176, and the pUC19-S-ecto-A-Nterm plasmid with primers MBK-177 and MBK-178. NEBuilder HiFi DNA Assembly protocol (NEB) was used to insert the kanamycin resistance gene into the plasmid. pUC19-S-ecto-Nterm was constructed by Golden Gate cloning (3) using pUC19-S-ecto-A-Nterm-KanR, pUC19-S-ecto-B, and pUC19-S-ecto-C-Nterm.

To construct pJS698 (N-terminal fusion **Spike ectodomain** YSD backbone), pETconNK-Nterm-Aga2p was first constructed by inserted a gene block with a multiple cloning site between the AGA2 signal peptide and the remainder of the AGA2 coding sequence following standard restriction enzyme cloning practices. pETconNK-Nterm-Aga2p was amplified with primers PJS-P2194 and PJS-P2195 using KAPA HiFi HotStart Readymix (Kapa Biosystems). The reaction was fractionated by agarose gel electrophoresis and the 6062 bp band excised and purified using a Monarch DNA Gel Extraction kit (NEB). The fragment (40 ng) was circularized using the Q5® Site-Directed Mutagenesis Kit (NEB) in a 10 µl reaction and transformed into E. coli Mach1 chemically competent cells (Invitrogen).

To construct pJS697 (C-terminal fusion **RBD** YSD backbone), pETconNK (Addgene: #81169) was amplified with primers PJS-P2192 and PJS-P2193 using KAPA HiFi HotStart Readymix (Kapa Biosystems). The reaction was fractionated by agarose gel electrophoresis and the 6084 bp band excised and purifed using a Monarch DNA Gel Extraction kit (NEB). The fragment (40 ng) was circularized using the Q5® Site-Directed Mutagenesis Kit (NEB) in a 10 µl reaction and transformed into *E. coli* Mach1 chemically competent cells (Invitrogen).

pJS699 (S-RBD(333-537)-N343Q for fusion to the C-terminus of AGA2) was previously described (4).

##### Recombinant protein production, purification, and preparation

ACE2-Fc was produced and purified following Walls et al. 2020 (5). CR3022 (6) was expressed by transient transfection in Expi293F cells and purified by protein A affinity chromatography and SEC using a Superdex 200 10/300 GL. Specificity was verified by measuring binding to SARS-CoV-2 RBD and irrelevant antigen. The anti-SARS-CoV-2 RBD antibody panel used (CC6.29, CC6.31, CC6.32, CC6.33, CC12.1, CC12.3, CC12.7, CC12.13, CC12.17, CC12.19) was a kind gift from Dennis Burton's lab at Scripps and were produced and purified according to Rogers et al. (7).

All proteins that were chemically biotinylated were prepared at a 20:1 molar ratio of biotin to protein using EZ-Link NHS-Biotin (Thermo Scientific) according to manufacturer's protocol. All proteins were stored at 4 °C in phosphate buffered saline (8 g/L NaCl, 0.2 g/L KCl, 1.44g/L $Na_2HPO_4$, 0.24g/L $KH_2PO_4$) pH 7.4.

**Preparation of Mutagenic Libraries**

All 119 surface exposed residues on S RBD (positions 333-537) were mutated to every other amino acid plus stop codon using NNK primers (**Table S5**) using comprehensive nicking mutagenesis exactly as described (8). For compatibility with Illumina sequencing, two tiles were made: tile 1 encompassed residues 333-436, while tile 2 encompassed residues 437-527 containing the critical receptor binding site. Serial dilutions were plated to calculate the transformation efficiency (**Table S1**). Glycerol stocks were made for each tile.

To create the display construct of S-RBD(333-537)-N343Q fused to the C-terminus of Aga2p, pJS697 was digested with BsaI-HFv2 (NEB) and purified using a Monarch PCR & DNA Cleanup Kit (NEB). Each mutated pJS699 library was digested with NotI-HF (NEB), the reaction fractionated by agarose gel electrophoresis, and the band corresponding to S-RBD (0.83kb) excised and purified using a Monarch DNA Gel Extraction Kit (NEB). Yeast transformation was performed exactly as described (9). For each library, the two fragments were co-transformed (in a 3:1 molar ratio of S-RBD to backbone) into chemically competent *S. cerevisiae* EBY100 (10). Serial dilutions were plated on SDCAA and incubated 3 days to calculate the efficiency of the transformation (**Table S1**). Biological replicates were made on a different day by co-transforming each tile into EBY100 exactly as described. Yeast stocks for each transformation were stored in yeast storage buffer at -80°C.

Mutagenic libraries for the N-terminal spike orientation were constructed following oligo pool mutagenesis exactly as described (8,11) using pUC19-S-ecto-A-Nterm-KanR, pUC19-S-ecto-B, and pUC19-S-ecto-C-Nterm as templates. For the oligo pool we computationally selected 1,909 mutations hypothesized to either destabilize the 'down' conformation, stabilize the 'up' conformation, or both (**Data S2**). The majority of these mutations targeted $S_1$ (94%, 1793/1909) at the NTD, RBD, SD1, and SD2 domains, with the remainder mapping to the boundary between the HR1 and CH domains on $S_2$. After mutagenesis, the mutational libraries were digested with BsaI-HFv2, fractionated by agarose gel electrophoresis, and gel excised and purified with Monarch Gel Extraction kit (NEB). 40 fmol of pUC19-S-ecto-A-NSM-Nterm-KanR, pUC19-S-ecto-B-NSM, and pUC19-S-ecto-C-NSM-Nterm were ligated together with T4 DNA Ligase (NEB), cleaned up and concentrated each to a final volume of 6µl with Monarch PCR & DNA Cleanup kit (NEB), and transformed into chemically competent *E.coli* Mach1 cells (Invitrogen cat. #C862003). The resulting library had on average 3 mutations per spike protein per plasmid. Library statistics were determined post sequencing.

To construct the surface display library in yeast, the spike plasmid library was digested with NotI-HF (NEB) and the S coding region was gel purified. The YSD vector pJS698 was digested with BsaI-HFv2 and column purified. 1.3 µg of insert (S coding region) and 1.7 µg of vector were electroporated into 400 μl EBY100 using the method of Benatuil et al. (12) as written, except that electroporation was performed at 2 kV rather than 2.5 kV. Immediately after electroporation, serial dilutions were plated on SDCAA Agar to calculate the complexity of the library. After electroporation, the cells were immediately transferred to 50 ml SDCAA (20g/L dextrose, 6.7g/L Difco yeast nitrogen base, 5g/L Bacto casamino acids, 5.4g/L $Na_2HPO_4$, and 8.56g/L $NaH_2PO_4 \cdot H_2O$ ) and grown at 30 °C for two days to saturation. The cultures were passaged twice in medium M37D (diluted to $OD_{600}$ = 0.05 in 120 ml, then to $OD_{600}$ = 0.4 in 50 ml) and stocks prepared at $OD_{600}$ = 1 as in Whitehead et al. (13). The final composition of M37 is 20 g $L^{-1}$ **d**extrose or **g**alactose (for M37D, M37G respectively), 5 g $L^{-1}$ casamino acids, 6.7 g $L^{-1}$ yeast nitrogen base with ammonium sulfate, 50 mM citric acid, 50 mM phosphoric acid, 80 mM MES acid, neutralized with 90% sodium hydroxide / 10% potassium hydroxide to pH 7. Both media should be prepared by dissolving all reagents except yeast nitrogen base into MilliQ water, adjusting the pH to 7.0 with freshly prepared sodium hydroxide / potassium hydroxide mixture, and adjusting the volume to 9/10$^{th}$ of the final desired volume. Pass the solution through a 0.22 μm filter, both for sterility and to remove particulates that would nucleate struvite. Finish the media by addition of 1/10$^{th}$ volume of 10x filtered yeast nitrogen base.

**Yeast Display Titrations and Competition Binding**

For cell surface titrations, EBY100 harboring the RBD display plasmid was grown in 1 ml M19D (5 g/l casamino acids, 40 g/l dextrose, 80 mM MES free acid, 50 mM citric acid, 50 mM phosphoric acid, 6.7 g/l yeast nitrogen base, adjusted to pH7 with 9M NaOH, 1M KOH) overnight at 30°C. Expression was induced by resuspending the M19D culture to $OD_{600}$=1 in M19G (5 g/l casamino acids, 40 g/l galactose, 80 mM MES free acid, 50 mM citric acid, 50 mM phosphoric acid, 6.7 g/l yeast nitrogen base, adjusted to pH7 with 9M NaOH, 1M KOH) and growing 22 h at 22°C with shaking at 300 rpm. For CR3022 IgG, yeast surface display titrations were performed as described by Chao et al. (14) with an incubation time of 4h at room temperature and using secondary labels anti-c-myc-FITC (Miltenyi Biotec) and Goat anti-Human IgG Fc PE conjugate (Invitrogen Catalog # 12-4998-82). Titrations were performed in biological replicates and technical triplicates (n = 6). The levels of display and binding were assessed by fluorescence measurements for FITC and SAPE using the Sony SH800 cell sorter equipped with a 70μm sorting chip and 488nm laser.

To test the individual antibody panel binding to S RBD, EBY100 harboring the RBD display plasmid was grown from -80°C cell stocks in 1 ml SDCAA for 4h at 30°C. Expression was induced by resuspending the SDCAA culture to $OD_{600}$=1 in SGCAA and growing at 22h at 22°C with shaking at 300rpm. 1x10$^5$ yeast cells were labelled with 10 $\mu$g/ml antibody IgG for 30min at room temperature in PBSF (PBS containing 1g/l BSA). The cells were centrifuged and washed with 200 μL PBSF. They were labeled with 0.6 μL FITC (Miltenyi Biotec), 0.25 μL Goat anti-Human IgG Fc PE (ThermoFisher Scientific) and 49.15 μL PBSF for 10min at 4°C. Cells were then centrifuged, washed with PBSF, and read on a flow cytometer to measure binding of the ACE2. Experiments were performed at least in biological replicate.

Competitive binding assays on the yeast surface were performed between a free antibody and biotinylated ACE2. *S. cerevisiae* EBY100 harboring the RBD display plasmid was grown from -80°C cell stocks in 1 ml SDCAA for 4h at 30°C. Expression was induced by resuspending the SDCAA culture to $OD_{600}$=1 in SGCAA and growing at 22h at 22°C with shaking at 300rpm. 1x10$^5$ yeast cells were labelled with 10 $\mu$g/ml antibody IgG for 30min at room temperature in PBSF (PBS containing 1g/l BSA). The same cells were labelled with 30nM chemically biotinylated hACE2, in the same tube without washing, for 30min at room temperature in PBSF. The cells were centrifuged and washed with 200 μL PBSF. They were labeled with 0.6 μL FITC (Miltenyi Biotec), 0.25 μL SAPE (Invitrogen) and 49.15 μL PBSF for 10min at 4°C. Cells were then centrifuged, washed with PBSF. The pellet was resuspended in 100 μL and read on a flow cytometer to measure binding of the hACE2.

**Yeast Display Screening of S and S RBD libraries**

For full-length S ectodomain screening, pUC19-S-ecto-Nterm and pJS698 were independently linearized via digest with restriction enzymes at 37°C for 1 hour, and gel extracted based off size using Monarch DNA Gel Extraction Kit. The linearized regions were co-transformed in a molar ratio of 1:3 insert to vector into chemically competent EBY100 following published protocols (9). EBY100 cells were recovered in nuclease free water for 5 minutes and then plated on two different yeast media agar plates: SDCAA and M37D. Cells were incubated at 30°C for 3 days. After initial growth, colonies from each plate were selected and grown up at 30°C and 250rpm overnight in the respective dextrose media: SDCAA, M37D. Cells were then induced in respective galactose media at an $OD_{600}$=1 at three different temperatures, 18°C, 22°C, and 30°C for 20 hours.

Induced EBY100 cells were washed with PBSF (8 g/L NaCl, 0.2 g/L KCl, 1.44g/L $Na_2HPO_4$, 0.24g/L $KH_2PO_4$, and 1g/L bovine serum albumin, pH to 7.4 and filter sterilized) and resuspended in PBSF at an $OD_{600}$=10. The cells were then incubated with either 500nM of the biotinylated ACE2-Fc or 500nM of the biotinylated CR3022 for 1 hour at room temperature. The cells were then washed with PBSF and labeled with anti-cmyc fluorescein isothiocyanate (FITC) (Miltenyi Biotec) and streptavidin phycoerythrin (SAPE) (Invitrogen) and incubated on ice for 10 minutes.

The Spike mutagenic library was labeled with CR3022 and, separately, ACE2-Fc under the optimal conditions were screened. Approximately $10^8$ were sorted using fluorescence activated cell sorting (FACS), and the top 1% of cells by fluorescence were collected. The two resulting sorted libraries were expanded and sorted in a second round, again screening $10^8$ cells and collecting the top 1% by fluorescence intensity. The selected populations were amplified and purified based on tile, deep sequenced, and count data compared with a reference population.

For the escape mutant screening of the S RBD, $3\times10^7$ induced EBY100 yeast cells displaying S RBD were labelled with 10 μg/ml antibody IgG for 30min at room temperature with mixing by pipetting every 10min in PBSF (PBS containing 1g/l BSA). The same cells were labelled with 75nM chemically biotinylated ACE2, in the same tube, for 30min at room temperature in PBSF with mixing by pipetting every 10min. The cells were centrifuged and washed with 1mL PBSF. Cells were then labeled with 1.2 μL FITC, 0.5 μL SAPE and 98.3 μL PBSF for 10min at 4°C. Cells were centrifuged, washed with 1mL PBSF, resuspended to 1 mL PBSF and sorted using FACS. Multiple gates were used for sorting as shown in **Figure S6**, including an $FSC/SSC^+$ gate for isolation of yeast cells, FSC-H/FSC-A gate to discriminate single cells, and the top 2% by a $PE^+/FITC^+$ gate. At least $3.0\times10^5$ cells were collected and were recovered in SDCAA with 50 μg/mL Kanamycin and 1x PenStrep for 30h. Biological replicates were sorted as described except for the ACE2 concentration being 30nM, maintaining saturation conditions.

**Deep Sequencing Preparation**

Libraries were prepared for deep sequencing following the "Method B" protocol from Kowalsky et al (15) exactly as described for the spike ectodomain libraries and with a few changes for the RBD libraries. A Monarch PCR & DNA Cleanup kit was used. PCR of extracted and cleaned-up yeast plasmid DNA was performed using 2xQ5 HotStart Master Mix (NEB) and the following protocol:

- 1 min @ 98 °C
- 25 cycles of:
  - 10 sec @ 98 °C
  - 20 sec @ 64 °C
  - 30 sec (replicate 1) or 1 min (replicate 2) @ 72 °C
- 2 min @ 72 °C
- Hold @ 4 °C

Primers used in library prep are given in **Table S5**. Amplicons were fractionated by agarose gel electrophoresis and purified using a Monarch DNA Gel Extraction Kit (NEB). Samples were then further purified using Agencourt Ampure XP beads (Beckman Coulter), quantified using PicoGreen (ThermoFisher), pooled, and sequenced on an Illumina MiSeq using 2 x 250 bp paired-end reads at the BioFrontiers Sequencing Core (University of Colorado, Boulder, CO).

**Deep Sequencing Analysis**

All deep sequencing data analysis was performed by scripts written in Python, available at GITHUB (https://github.com/WhiteheadGroup/SpikeRBDStabilization.git).

Because all sequenced samples were PCR amplicons of known length, paired-end reads were merged by aligning at the known overlap. Mismatches in overlapping regions were resolved by selecting the base pair with the higher quality score and assigning it a quality score given by the absolute difference of the quality scores at the mismatch. Paired reads with more than 10 mismatches in the overlapping region and merged reads containing any quality score less than 10 were discarded. The total number of retained reads in each sample was recorded as $n_i$, the number of reads in sample $i$.

Each read was compared to the wild-type sequence to identify all mutations. Counts for synonymous single mutations were combined to give $k_{ij}$, the number of reads in sample $i$ encoding the single amino acid mutation $j$. Reads including multiple mutations or mutations not encoded in the library oligos were not analyzed further. The frequency of single mutant $j$ in sample $i$ was calculated as $f_{ij} = k_{ij} / n_i$.

Each experiment consisted of two samples: a reference sample $r$ and a selected sample $s$. For each experiment, the risk ratio of variant $j$ was calculated as $\rho_j = f_{sj} / f_{rj}$ i.e. the ratio of the variant's frequency in the selected population to its frequency in the reference population. Enrichment ratios were calculated as the binary logarithm of the risk ratio: $ER_j = \rho_j$ . Variants with five or fewer counts in the reference population were not analyzed further. Variants with at least five counts in the reference population but no counts in the selected population were given a pseudocount of one.

*Determining hits from yeast display screens*

For each escape mutant screen, we collected the top 2% (PE channel) of the population of FITC$^{+}$ (RBD displaying) cells. This population was not labeled with biotinylated ACE2 and so serves as a null experiment where the observed enrichment ratios are due to other sources of variance and not to differential nAb binding. We fit the distribution of enrichment ratios for each of these control samples using kernel density estimation (KDE) (SciPy's scipy.stats.gaussian_kde with default parameters) (16). We then treated this distribution estimate as an empirical null hypothesis. Under this null hypothesis, we expect $N\big(1 - F(ER_t)\big)$ false positives, where N is the number of variants tested, F is the cumulative distribution function (CDF) of the control ER KDE, and $ER_t$ is a threshold. Therefore, for a target false discovery rate (FDR), we chose $ER_t = F^{-1}(1 - FDR/N)$, where $F^{-1}$ is the inverse CDF of the KDE. In data from samples labeled with nAbs, we then tested the hypothesis that each observed ER was greater than the associated $ER_t$ using an one-sided exact Poisson rate ratio test (statsmodels.stats.rates.test_poisson_2indep from the Python library statsmodels) (17). For these tests, the null ratio was $2^{ER_t}$. The counts were given by the number of reads for the variant in the selected and reference populations, respectively, and the exposures were given by the total number of reads in the reference and selected populations, respectively. For this analysis, we identified hits for replicate 1 (tiles 1 & 2 for nAbs CC6.29, CC12.1, and CC12.3) using a target FDR of 1 and a Poisson rate ratio test significance level of 0.01. For replicate 2 (tile 2 for nAbs CC6.31, CC12.13) escape mutant hits were identified using a target FDR of 1.

For the full-length S ectodomain screen, our null experiment was the collected reference populations without selections for each of the ACE2-Fc and CR3022 experiments. These reference populations were passaged, sorted, and amplified identically to the sorted libraries except that no screen was employed. We fit the distribution of enrichment ratios for these control samples using a logistic CDF (custom MATLAB script), and the empirical FDR was calculated exactly as above.

**Molecular Dynamics Simulations**

GROMACS 2018.3 (25) was employed for all molecular dynamics (MD) simulations along with the TIP3P (26) water model and Amber99SB-ILDN (27) force field to model the receptor binding domain (RBD) of the spike (S) protein of SARS-CoV-2 and neutralizing antibodies CC12.1 and CC12.3. Simulations were initiated from crystal structures of the RBD in complex with CC12.1 (PDB code 6XC2 (28)) and CC12.3 (PDB code 6XC4 (28)). All systems containing a positive charge were neutralized by the addition of Cl- ions, also modeled with the Amber99SB-ILDN force field. Each simulation consisted of approximately 192,000 atoms.

A steepest descent energy minimization of the initial coordinates for each system was carried out for 5,000 steps. NVT equilibration simulations were then performed for 0.5 ns at 310 K with the Bussi−Donadio−Parrinello (29) thermostat. Subsequent NPT equilibration simulations were performed for 1 ns at 310 K and 1.0 bar, using the same thermostat and Berendsen (30) barostat. The time constant for coupling in both the NVT and NPT simulations was 0.1 ps. Production simulations in the NPT ensemble were then carried out at 310 K and 1.0 bar with the Bussi−Donadio−Parrinello thermostat and Parrinello–Rahman (31) barostat. Long-range electrostatic interactions were calculated using particle mesh Ewald summations and a cutoff of 1.0 nm, and Lennard Jones interactions were calculated over 1.0 nm and shifted beyond this distance. Neighbor lists were updated every 10 steps with a cutoff of 1.0 nm. Bonds between hydrogen and heavy atoms were constrained with the LINCS (32) algorithm. Furthermore, periodic boundary conditions were used in all simulations in all directions. Production simulations were carried out for 100 ns, leading to a total of 0.8 microseconds of simulation time across the eight simulations.

**Pseudo Neutralization Assays**

SARS-CoV-2 pseudovirus neutralization assays were performed as previously described (7). Briefly, pseudovirus was generated by cotransfecting MLV-gag/pol and MLV-CMV-Luciferase plasmids with truncated wildtype SARS-CoV-2 or mutant SARS-CoV-2 plasmid respectively onto HEK293T cells. After 48h or 72h of transfection, supernatants containing pseudovirus were collected and frozen at -80 °C. Neutralization assay was performed as follows. First, monoclonal antibodies were serially diluted into half-area 96-well plates (Corning, 3688) and incubated with pseudovirus at 37 °C for 1 h. Next, HeLa-hACE2 cells were transferred in the 96-well plates at 10,000 cells/well. After 48h of incubation, supernatants were removed, cells were lysed with 1x luciferase lysis buffer (25 mM Gly-Gly pH 7.8, 15 mM $MgSO_4$, 4 mM EGTA, 1% Triton X-100). Finally, Bright-Glo (Promega, PR-E2620) was added onto 96-well plates according to manufacturer's instructions. Neutralization IC50s were calculated using "One-Site LogIC50" regression in GraphPad Prism 8.0. Pseudovirus mutant constructs were generated by amplifying two overlapped fragments of SARS-CoV-2 mutant sequences with Q5 enzyme (NEB, M0492) following manufacturer's instructions. Two fragments were then joint into one fragment by bridge PCR,  and gibson cloned into digested pcDNA3.3 backbone.

#### FIGURES

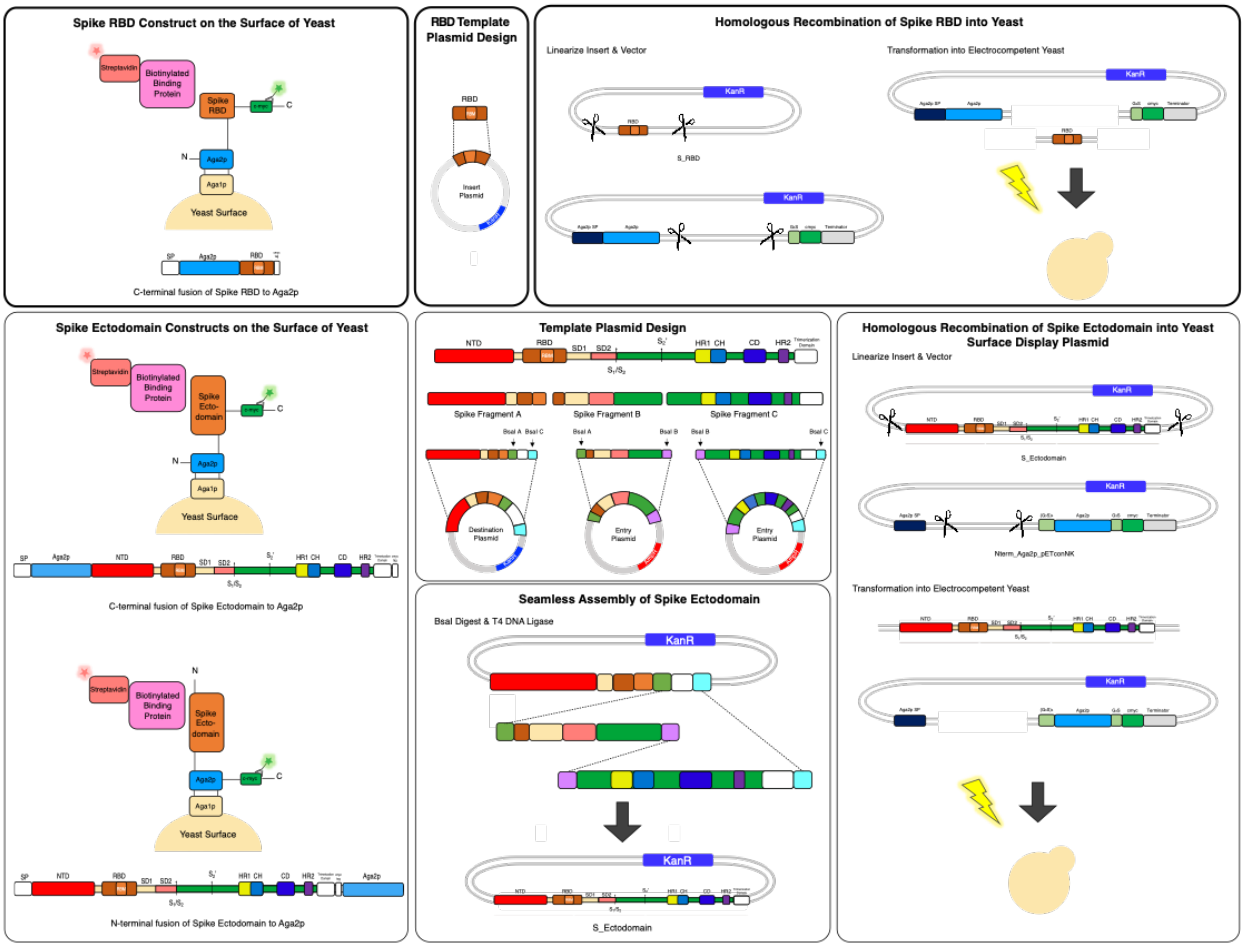


**Fig. S1. Overview of yeast display constructs used in screening S RBD and prefusion stabilized S ectodomain libraries.**

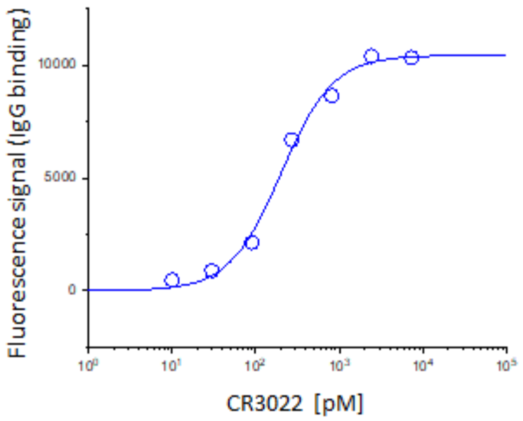


| IgG | Apparent $K_d$, YSD [pM] | Apparent $K_d$, *in vitro* [pM] | Reference |
|---|---|---|---|
| CR3022 | 171±25 | ~ 100 | Yuan, et al. 2020 |
| 910-30 | 230±70 | 104 | Banach, et al. 2021 |

**Fig. S2. Non-neutralizing antibody CR3022 dissociation constant compared to literature.** Yeast cell surface titrations of non neutralizing CR3022 IgG against aglycosylated S RBD yield an apparent $K_D$ of 171 ± 25 pM. Technical triplicates were performed (n = 3), and error reported is 2 s.e.m. Data for the HKU 910-30 nAb is from Banach et al, 2021. (4,18).

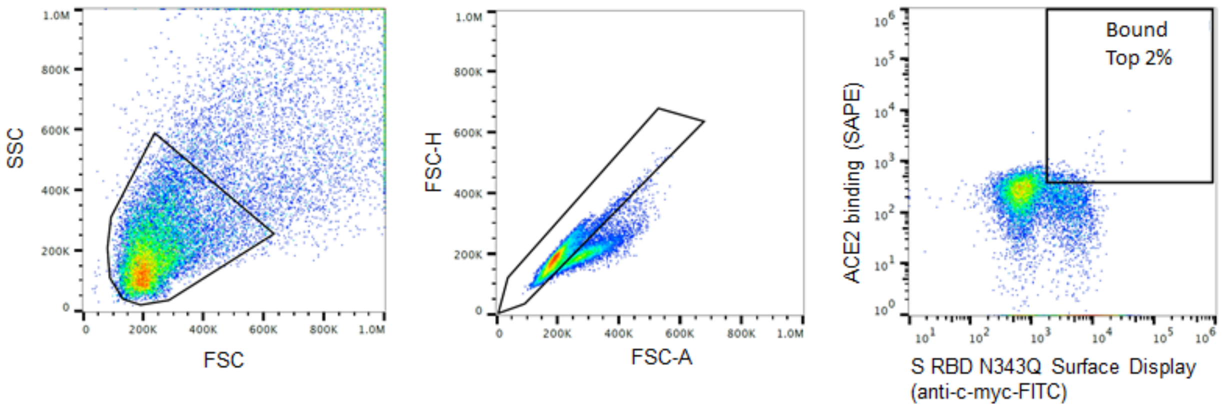


**Fig. S3. FACS sorting gates used to collect S RBD escape mutants**
Representative sorting gates used for all escape mutant FACS screens. The three gates were SSC/FSC; FSC-H/FSC-A to discriminate single yeast cells; and $SAPE^+/FITC^+$ to identify mutants that allow ACE2 binding in the presence of 10 $\mu$g/mL antibody. Shown here are gates for the antibody CC6.29.

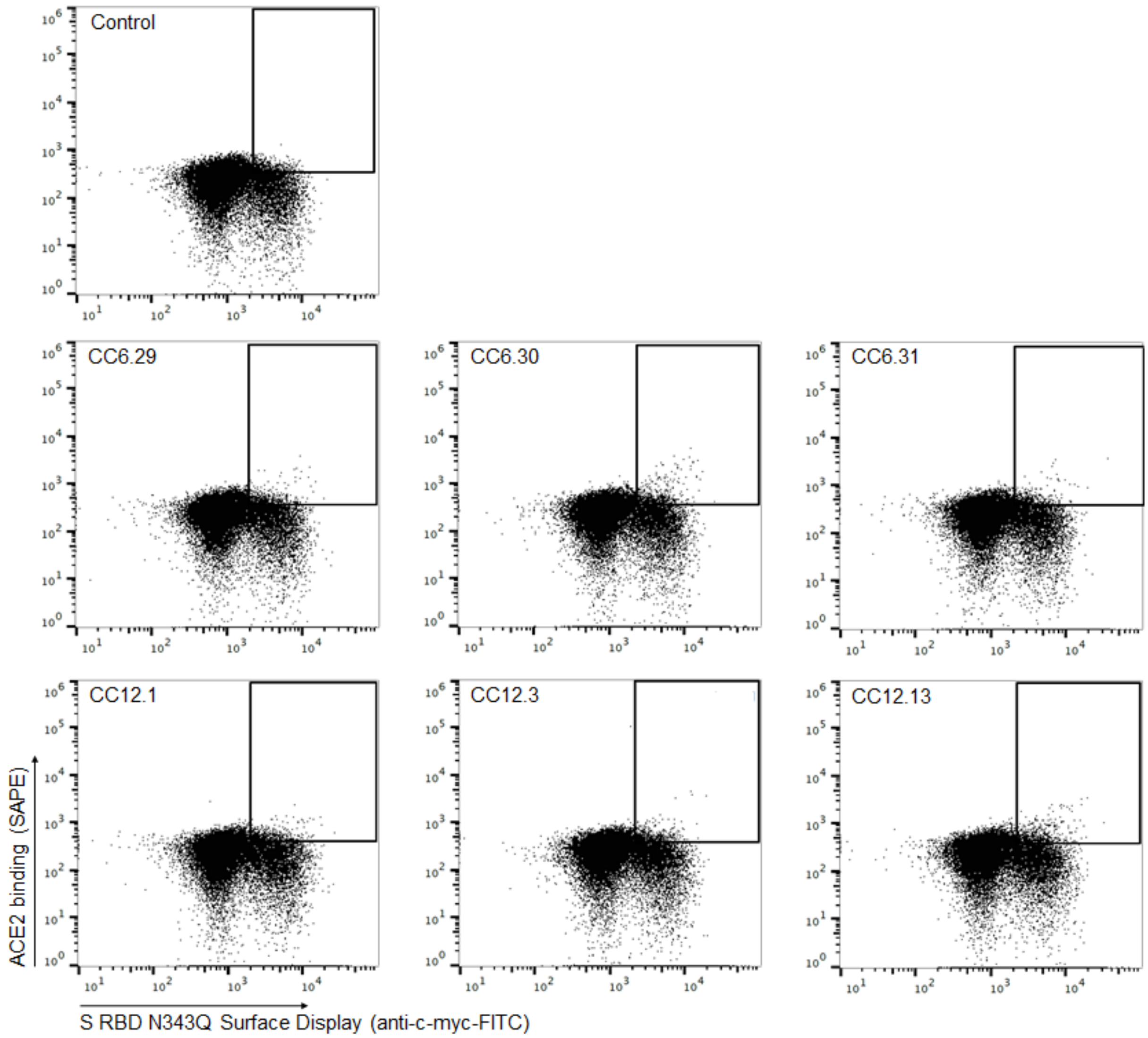


**Fig. S4. Sorting gates for competitive binding experiments.**
PE/FITC cytograms for aglycosylated S RBD yeast libraries sorted using competitive binding with ACE2-Fc. The top cytogram shows the control experiment with no ACE2 labeling. Gates represent the top 2% of the $FITC^+$ cells by PE signal for each antibody used in the study.

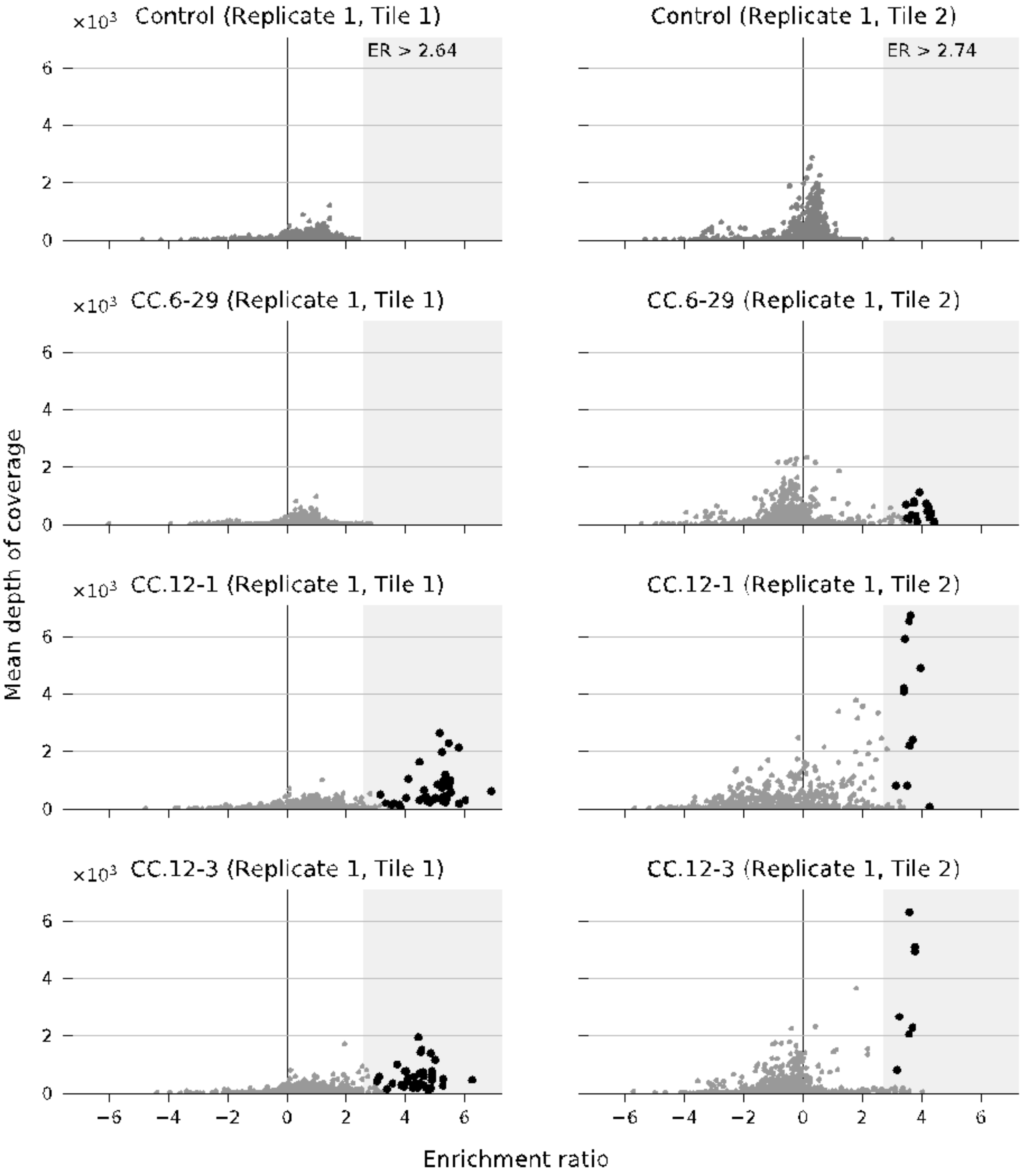


**Fig. S5.** Per-mutation enrichment ratio (ER) distributions as a function of average depth of coverage control (top) and CC12.3, CC12.1, CC6.29 nAb competing experiment (bottom). ER thresholds determined for FDR = 1 are shown in the control panels (top). Hits (with ER greater than the threshold at p <= 0.01) are shown with larger black dots.

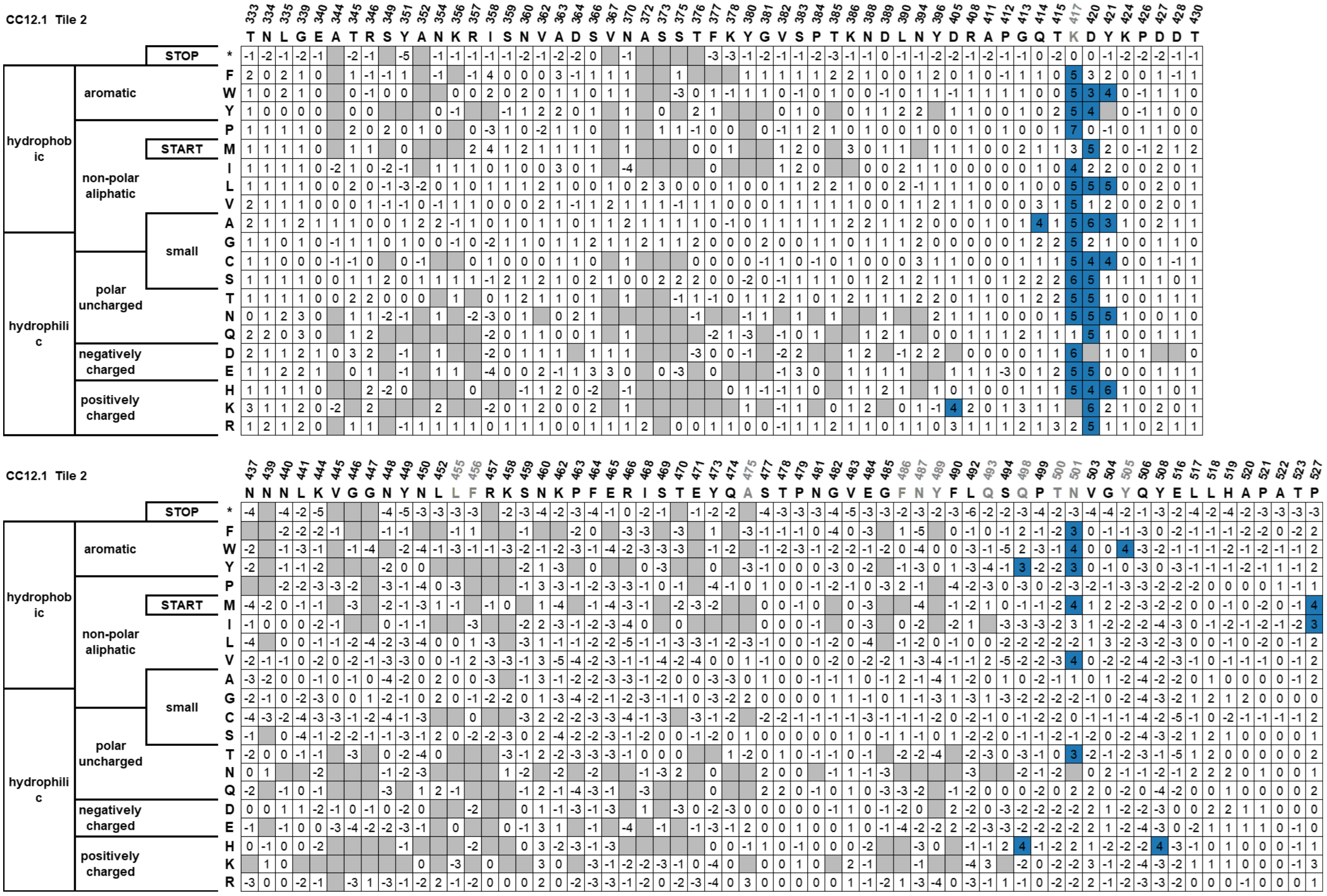


**Fig. S6.** CC12.1 heatmap using FDR < 0.1 as cut-off. Blue - escape mutant hit, white – not a hit, grey – mutation not present. Position numbers in grey indicate ACE2 footprint.

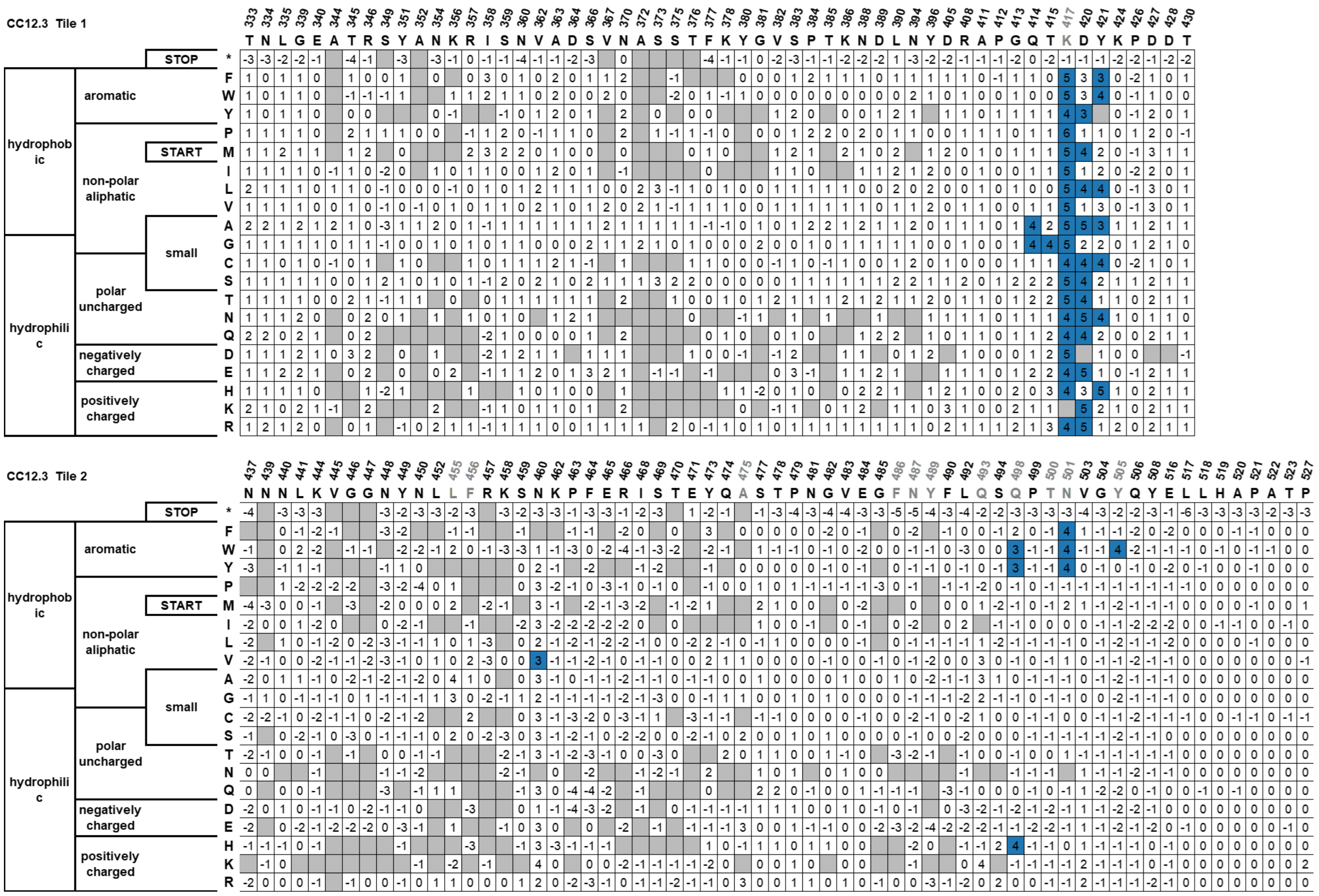


**Fig. S7.** CC12.3 heatmap using FDR < 0.1 as cut-off. Blue - escape mutant hit, white – not a hit, grey – mutation not present. Position numbers in grey indicate ACE2 footprint.

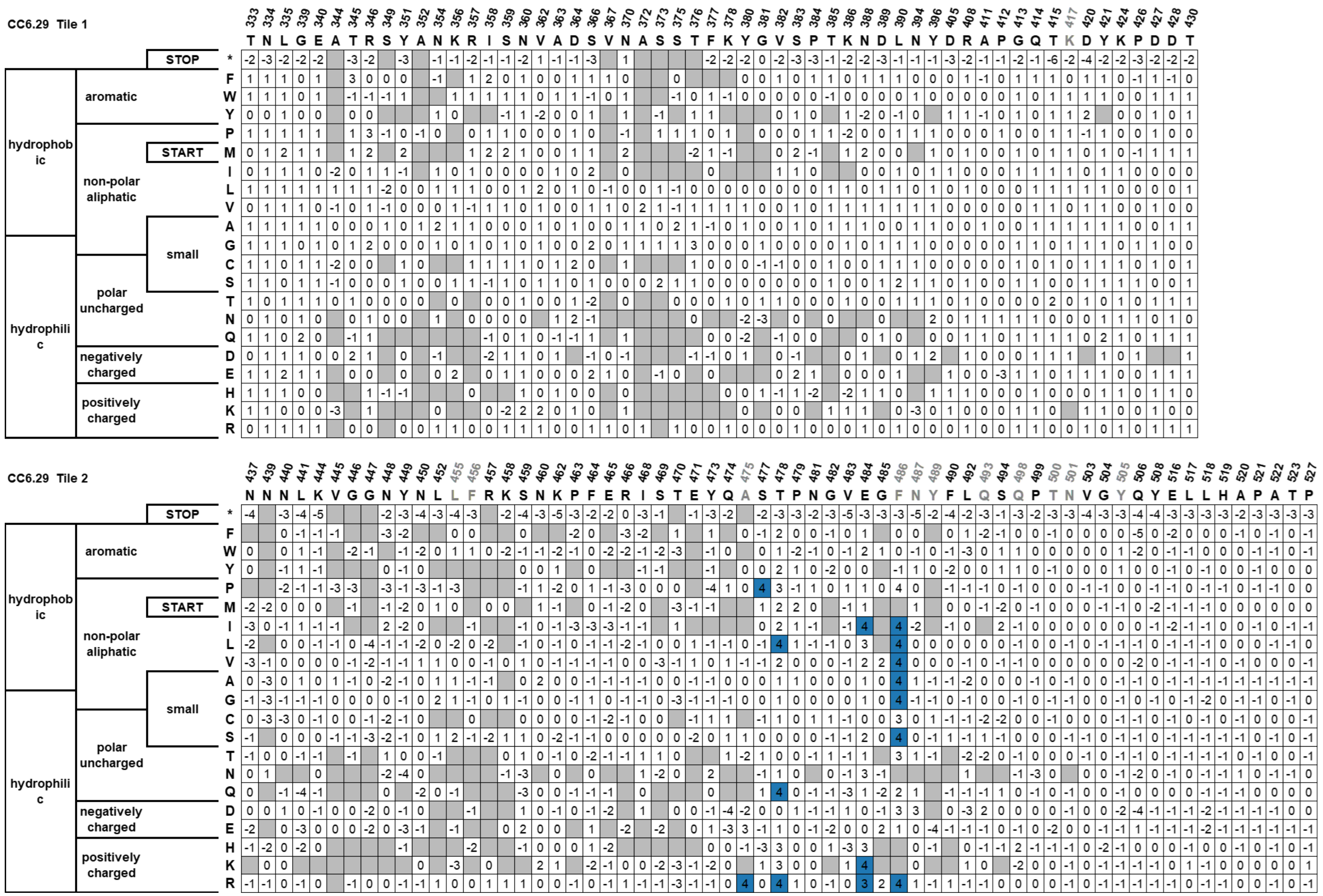


**Fig. S8.** CC6.29 heatmap using FDR < 0.1 as cut-off. Blue - escape mutant hit, white – not a hit, grey – mutation not present. Position numbers in grey indicate ACE2 footprint.

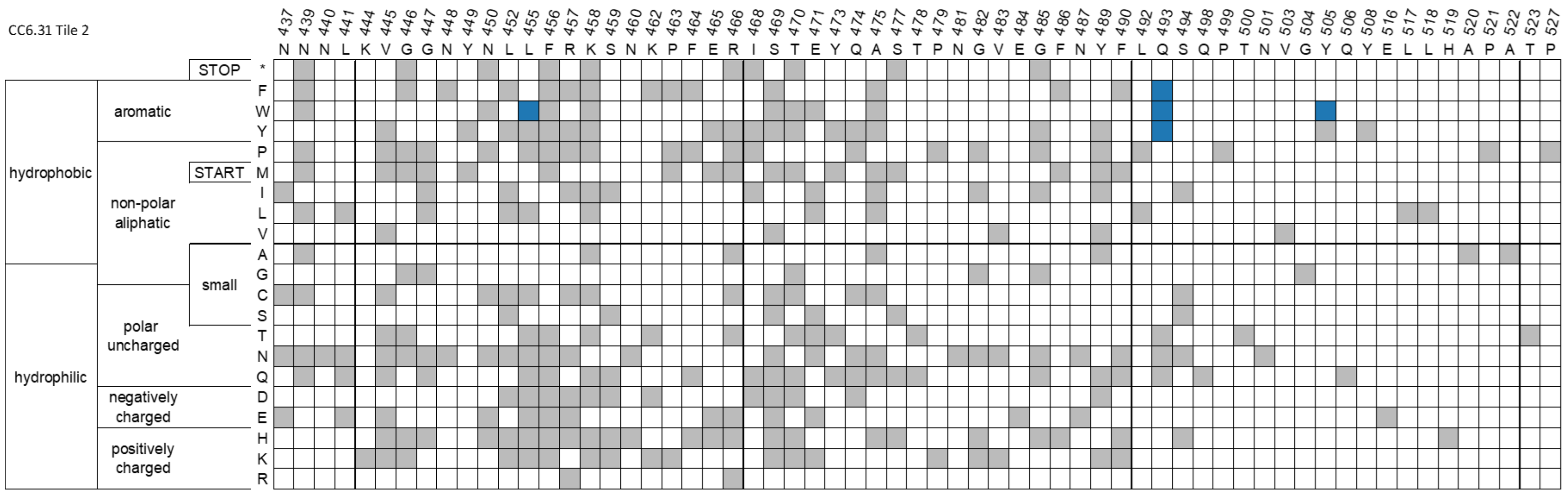


**Fig. S9.** CC6.31 heatmap using FDR < 1 as cut-off. Blue - escape mutant hit, white – not a hit, grey – mutation not present. Position numbers in grey indicate ACE2 footprint.

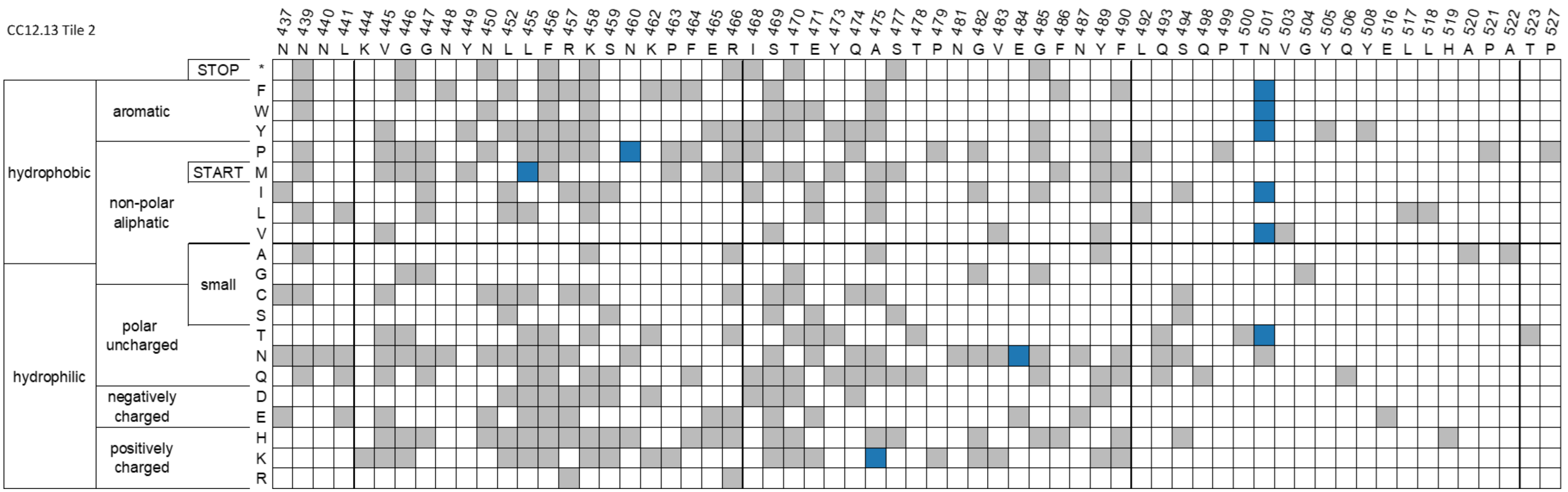


**Fig. S10.** CC12.13 heatmap using FDR < 1 as cut-off. Blue - escape mutant hit, white – not a hit, grey – mutation not present. Position numbers in grey indicate ACE2 footprint.

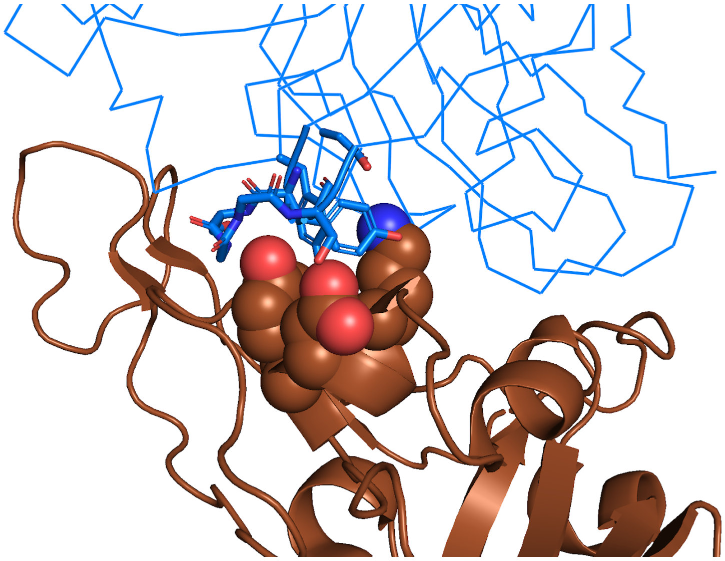

**Fig. S11.** Structural recognition of S RBD (chocolate cartoon) by nAB CC12.1 (PDB ID 6XC2, blue ribbon). S positions K417, D420, Y421 are shown as spheres and the CDR H2 and key CC12.1 residues are shown as sticks.

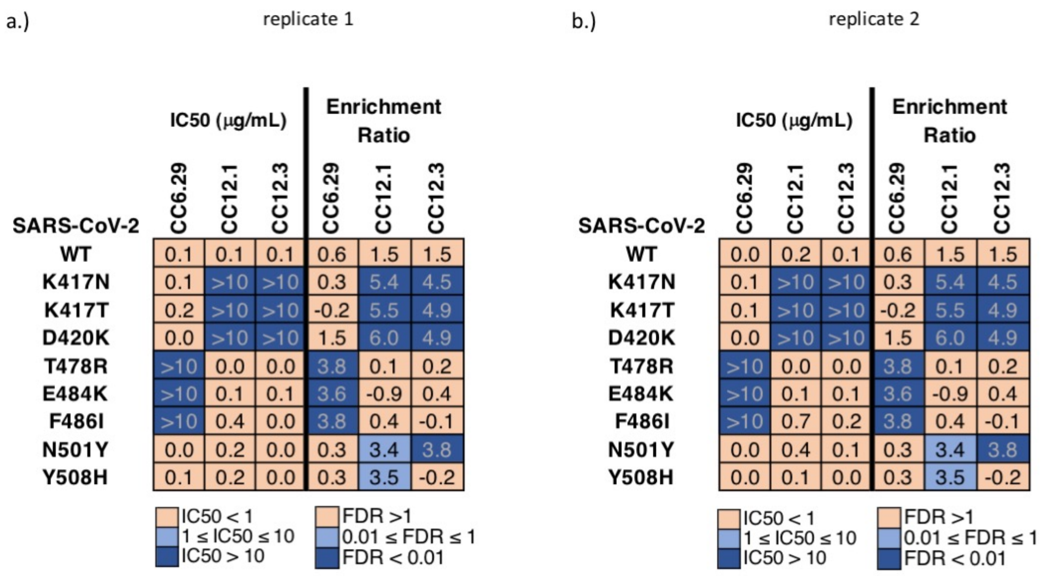


**Fig. S12.** MLV-based SARS-CoV-2 pseudovirus neutralization assays for SARS-CoV-2 RBD variants. Data shown are replicates of the neutralization assays repeated on separate days (replicate 1 - day 1; replicate 2 - day 2). IC50 neutralization values shown are averages of two technical replicates. Replicate 1 data is also presented in Figure 1j of the main text.

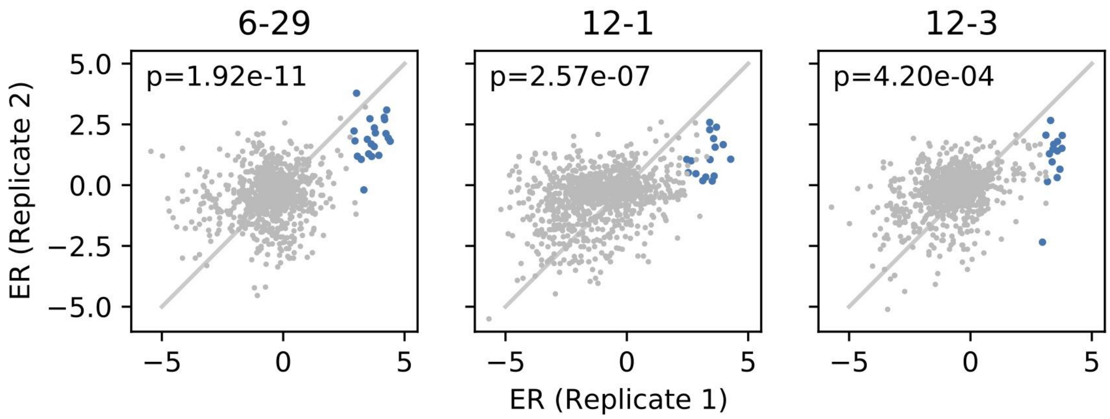


**Fig. S13. Comparison between biological replicates for S RBD positions 437-527 for CC6.29, CC12.1, and CC12.3.** Escape mutant hits identified in replicate 1 are shown as closed blue circles (a p <= 0.01 for an FDR < 1). p-values are calculated using a one-sided Welch's t-test with the alternative hypothesis that the mean enrichment ratio from the replicate 2 hits are > the mean enrichment ratio from the replicate 2 non-hits.

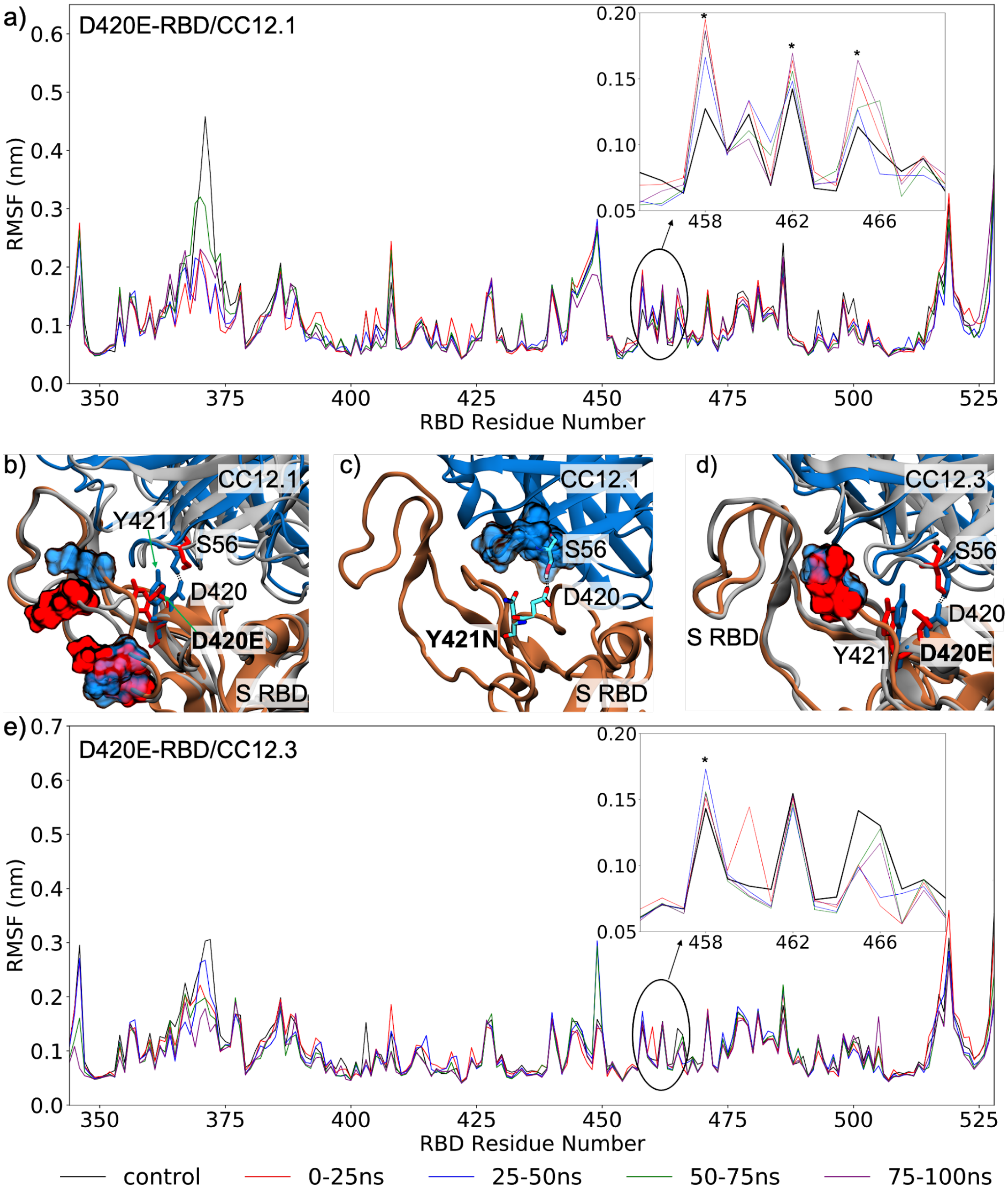


**Fig. S14.** (a) Comparison of the root mean square fluctuation (RMSF) profile of CC12.1 in complex with wildtype/control S RBD (averaged across the 100 ns production run) and S RBD with the D420E mutation (averaged across 25 ns intervals). (b) Structural mapping of highly fluctuating residues on S RBD with the D420E mutation, identified in panel (a), when complexed with CC12.1. Wildtype and mutant RBDs are shown in brown and gray and CC12.1 in blue and gray, respectively, while residues are colored blue for wildtype RBD and red for the mutant RBD; highly fluctuating residues are shown in surface representation. (c) MD snapshot showing the proposed mechanism of escape of S RBD from CC12.1 through mutation Y421N. (d) Structural mapping of highly fluctuating residues on S RBD with the D420E mutation, identified in panel (e), when complexed with CC12.3. (e) Comparison of the RMSF profile of CC12.3 in complex with wildtype S RBD and S RBD with the D420E mutation.

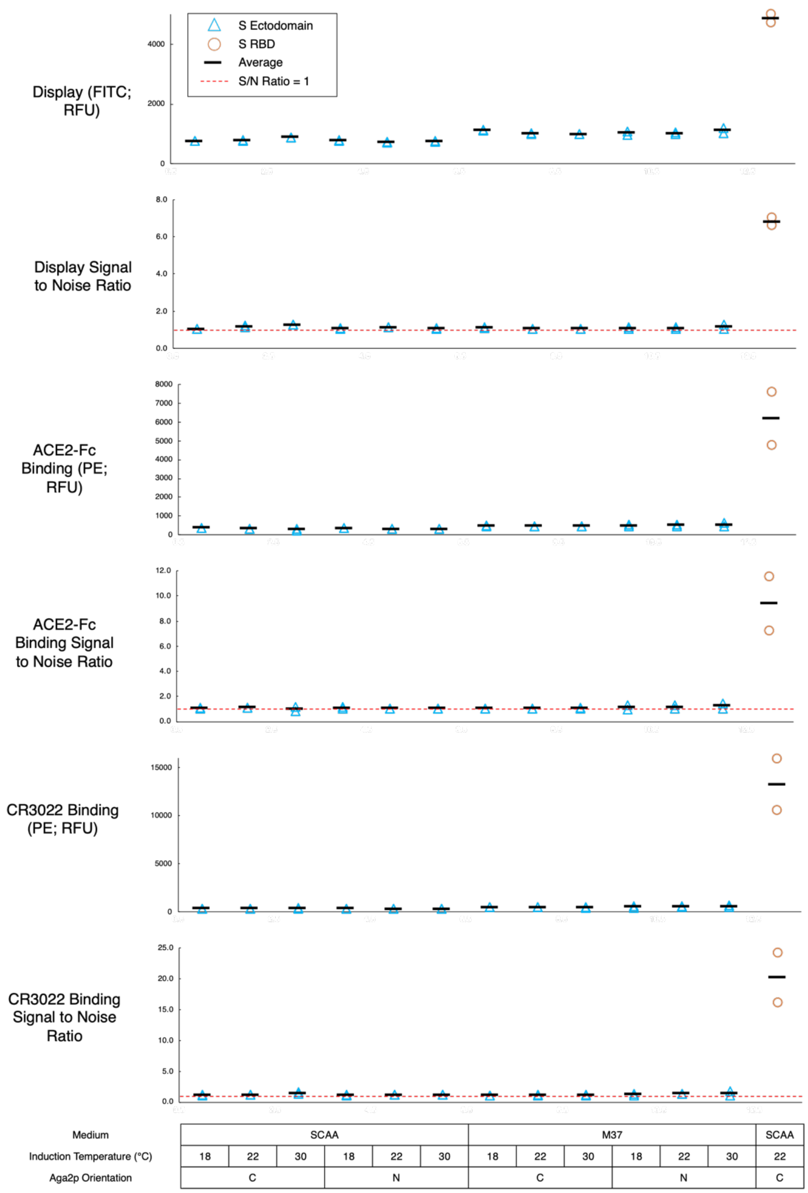


**Fig. S15: Yeast surface display for SARS-CoV-2 prefusion stabilized S ectodomain compared to S RBD**. FITC signal (RFU), FITC signal to noise ratio, and PE signal (RFU) and PE signal to noise ratio for both ACE2-Fc and CR3022 are shown for biological replicates of spike ectodomain with differing media, induction temperatures, and orientation of Spike relative to Aga2p (blue triangles). The FITC fluorescence derives from an anti-cmyc FITC antibody that recognizes a C-terminal cmyc epitope tag for displayed protein, while the PE signal is from biotinylated ACE2-Fc or CR3022 subsequently labeled with streptavidin-PE. The concentration of the secondary binding protein for the S ectodomain was 500nM ACE2-Fc and 500nM CR3022. FITC signal (RFU), FITC signal to noise ratio, and PE signal (RFU) and PE signal to noise ratio for both ACE2-Fc and CR3022 shown for S RBD at optimal conditions (orange circles). The concentration of secondary binding protein for the S RBD is at 1 nM, which is at saturation.

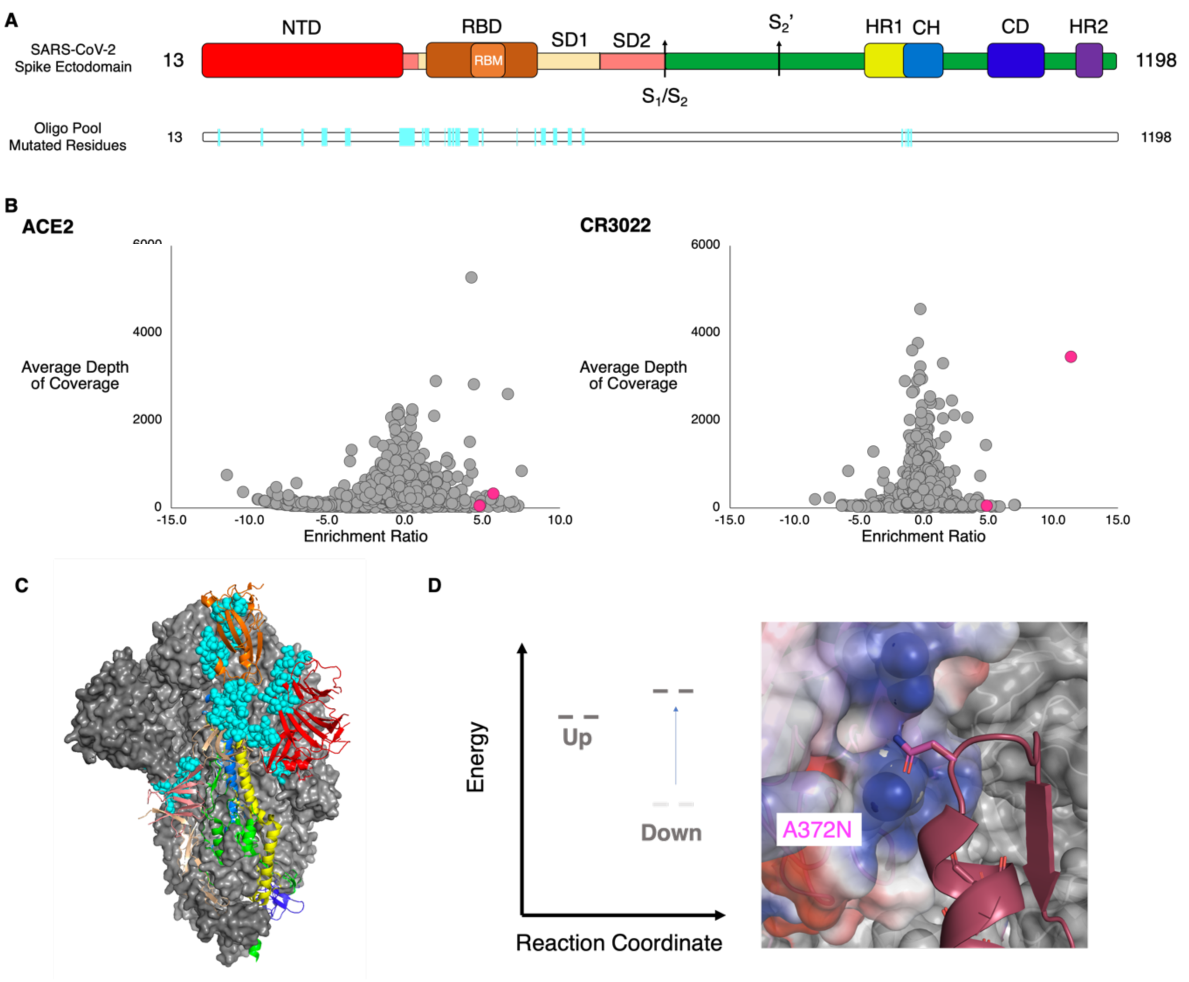


**Fig. S16. Determination and location of potential stabilizing hits.** (A) Spike ectodomain schematic with labeled and colored boundaries. Below schematic is the locations of the mutated residues in the oligo pool, cyan, as well as locations of the top identified hits shown in magenta. NTD: N-terminal domain, RBD: receptor-binding domain, RBM: receptor-binding motif, SD1: subdomain 1, SD2: subdomain 2, $S_1/S_2$: furin cleavage site, $S_2$': $S_2$' cleavage site, HR1: heptad repeat 1, CH: central helix, CD: connector domain, HR2: heptad repeat 2. (B) Average depth of coverage vs. Enrichment ratio for single mutants for (left) ACE2 and (right) CR3022. All mutations are shown in gray with the two hits (K113I & A372N) colored magenta. (C) Prefusion spike trimer shown with domains colored as they are in panel A. RBD is shown in the up conformation and along with SD1 and SD2 is shown on the same spike monomer. The NTD and S2 subunits are shown on a neighboring monomer. Oligo pool mutated residues are represented as cyan spheres. (D) Putative reaction coordinate and interaction of hit A372N in the spike ectodomain. A372N is hypothesized to destabilize the down protomer by steric repulsion with an adjacent 'down' protomer.

**TABLES**
**Table S1.**
Summary of Statistics for S RBD & N-Term Spike ectodomain Libraries. Library statistics for S RBD were determined by tabulating all variants with at least two reads in the yeast reference libraries. Library statistics for S ectodomain were determined from NGS of the libraries harbored in *E. coli*.

| | **S RBD** | | **S Ecto** | | | | |
|---|---|---|---|---|---|---|---|
| Tile Number | Tile 1 | Tile 2 | Tile 1 | Tile 2 | Tile 3 | Tile 4 | Tile 5 |
| Positions | 333-436 | 437-537 | 31-177 | 189-341 | 355-494 | 510-589 | 936-1020 |
| Number of Designed Mutations | 1120 | 1260 | 229 | 582 | 639 | 324 | 135 |
| Transformants Obtained from Nicking Saturation Mutagenesis | 1.10E+06 | 2.70E+07 | | 2.00E+06 | | 1.00E+06 | 1.00E+04 |
| Transformants Obtained from Assembly of Spike Ectodomain | N/A | | | | 4.00E+05 | | |
| Transformants Obtained from Homologous Recombination | 2.50E+05 | 1.10E+05 | | | 1.75E+06 | | |
| Library Coverage Per Tile | 97% (1085/1120) | 94% (1189/1260) | 88% (201/229) | 90% (523/582) | 90% (577/639) | 100% (324/324) | 99% (134/135) |
| Overall Library Coverage | 96% (2274/2380) | | | | 92% (1759/1909) | | |

**Table S2**. List of escape mutants identified in this study. Escape mutants were identified by a p value < 0.01 for containing a FDR < 1.

| nAb | Variant | Counts in competition selection | Counts in reference population | Enrichment Ratio | Minimum nucleotide distance | FDR | p value for FDR < 1 |
|---|---|---|---|---|---|---|---|
| CC.12-1 | D405K | 249 | 24 | 3.5 | 2 | 7.7E-07 | 1.5E-03 |
| CC.12-1 | Q414A | 566 | 28 | 4.5 | 2 | 5.0E-13 | 8.3E-16 |
| CC.12-1 | K417A | 634 | 19 | 5.2 | 2 | 5.0E-13 | 6.8E-25 |
| CC.12-1 | K417C | 506 | 21 | 4.7 | 3 | 5.0E-13 | 1.9E-16 |
| CC.12-1 | K417D | 358 | 7 | 5.8 | 2 | 5.0E-13 | 1.4E-17 |
| CC.12-1 | K417E | 800 | 23 | 5.3 | 1 | 5.0E-13 | 1.0E-31 |
| CC.12-1 | K417F | 446 | 12 | 5.4 | 3 | 5.0E-13 | 6.3E-19 |
| CC.12-1 | K417G | 1655 | 54 | 5.1 | 2 | 5.0E-13 | 1.9E-59 |
| CC.12-1 | K417H | 432 | 17 | 4.8 | 2 | 5.0E-13 | 8.5E-15 |
| CC.12-1 | K417I | 155 | 12 | 3.8 | 1 | 1.3E-10 | 1.6E-03 |
| CC.12-1 | K417L | 2303 | 63 | 5.3 | 2 | 5.0E-13 | 1.1E-90 |
| CC.12-1 | K417N | 732 | 19 | 5.4 | 1 | 5.0E-13 | 1.1E-30 |
| CC.12-1 | K417P | 1197 | 11 | 6.9 | 2 | 5.0E-13 | 3.0E-68 |
| CC.12-1 | K417S | 1869 | 45 | 5.5 | 2 | 5.0E-13 | 1.1E-78 |

| | | | | | | | |
|---|---|---|---|---|---|---|---|
| CC.12-1 | K417T | 1528 | 38 | 5.5 | 1 | 5.0E-13 | 1.2E-63 |
| CC.12-1 | K417V | 787 | 33 | 4.7 | 2 | 5.0E-13 | 1.5E-24 |
| CC.12-1 | K417W | 1427 | 42 | 5.2 | 2 | 5.0E-13 | 1.2E-54 |
| CC.12-1 | K417Y | 523 | 15 | 5.3 | 2 | 5.0E-13 | 2.8E-21 |
| CC.12-1 | D420A | 4199 | 84 | 5.8 | 1 | 5.0E-13 | 5.5E-189 |
| CC.12-1 | D420C | 707 | 48 | 4.0 | 2 | 7.5E-13 | 1.1E-13 |
| CC.12-1 | D420E | 3842 | 113 | 5.2 | 1 | 5.0E-13 | 2.7E-144 |
| CC.12-1 | D420H | 372 | 34 | 3.6 | 1 | 1.1E-07 | 3.9E-05 |
| CC.12-1 | D420K | 584 | 10 | 6.0 | 2 | 5.0E-13 | 4.0E-29 |
| CC.12-1 | D420L | 1885 | 54 | 5.3 | 2 | 5.0E-13 | 9.8E-73 |
| CC.12-1 | D420M | 756 | 26 | 5.0 | 3 | 5.0E-13 | 3.9E-27 |
| CC.12-1 | D420N | 914 | 24 | 5.4 | 1 | 5.0E-13 | 1.0E-37 |
| CC.12-1 | D420Q | 678 | 24 | 5.0 | 2 | 5.0E-13 | 4.8E-24 |
| CC.12-1 | D420R | 4431 | 110 | 5.5 | 2 | 5.0E-13 | 1.8E-181 |
| CC.12-1 | D420S | 1979 | 49 | 5.5 | 2 | 5.0E-13 | 4.0E-82 |
| CC.12-1 | D420T | 733 | 33 | 4.6 | 2 | 5.0E-13 | 1.0E-21 |
| CC.12-1 | D420W | 372 | 41 | 3.3 | 3 | 5.8E-05 | 1.5E-03 |
| CC.12-1 | D420Y | 299 | 24 | 3.8 | 1 | 6.4E-10 | 2.4E-05 |

| | | | | | | | |
|---|---|---|---|---|---|---|---|
| CC.12-1 | Y421A | 888 | 109 | 3.2 | 2 | 1.1E-03 | 9.5E-05 |
| CC.12-1 | Y421C | 1952 | 126 | 4.1 | 1 | 5.0E-13 | 1.9E-37 |
| CC.12-1 | Y421H | 1140 | 27 | 5.5 | 1 | 5.0E-13 | 5.6E-49 |
| CC.12-1 | Y421L | 5137 | 159 | 5.2 | 2 | 5.0E-13 | 1.1E-186 |
| CC.12-1 | Y421N | 1262 | 56 | 4.6 | 1 | 5.0E-13 | 1.1E-36 |
| CC.12-1 | Y421W | 3129 | 156 | 4.5 | 2 | 5.0E-13 | 1.9E-79 |
| CC.12-1 | Q498H | 4086 | 296 | 3.6 | 1 | 1.7E-02 | 6.9E-30 |
| CC.12-1 | Q498Y | 1467 | 147 | 3.1 | 2 | 3.7E-01 | 2.0E-04 |
| CC.12-1 | N501F | 7508 | 617 | 3.4 | 2 | 7.6E-02 | 3.0E-37 |
| CC.12-1 | N501M | 9286 | 519 | 4.0 | 2 | 2.0E-04 | 4.0E-111 |
| CC.12-1 | N501T | 10942 | 890 | 3.4 | 1 | 6.8E-02 | 3.1E-55 |
| CC.12-1 | N501V | 12572 | 885 | 3.6 | 2 | 1.1E-02 | 1.7E-94 |
| CC.12-1 | N501W | 4503 | 305 | 3.7 | 3 | 6.3E-03 | 2.9E-38 |
| CC.12-1 | N501Y | 7767 | 639 | 3.4 | 1 | 7.6E-02 | 2.5E-38 |
| CC.12-1 | Y505W | 12149 | 894 | 3.6 | 2 | 2.1E-02 | 9.4E-82 |
| CC.12-1 | Y508H | 1478 | 113 | 3.5 | 1 | 3.3E-02 | 1.8E-10 |
| CC.12-1 | P527I | 467 | 43 | 3.3 | 2 | 2.1E-01 | 7.4E-03 |
| CC.12-1 | P527M | 155 | 7 | 4.3 | 2 | 1.3E-06 | 7.5E-04 |

| | | | | | | | |
|---|---|---|---|---|---|---|---|
| CC.12-3 | Q414A | 627 | 28 | 4.4 | 2 | 5.0E-13 | 1.1E-13 |
| CC.12-3 | Q414G | 807 | 44 | 4.1 | 2 | 5.0E-13 | 9.8E-14 |
| CC.12-3 | T415G | 1424 | 82 | 4.0 | 2 | 1.4E-12 | 2.3E-21 |
| CC.12-3 | K417A | 514 | 19 | 4.6 | 2 | 5.0E-13 | 1.7E-13 |
| CC.12-3 | K417C | 496 | 21 | 4.4 | 3 | 5.0E-13 | 1.1E-11 |
| CC.12-3 | K417D | 210 | 7 | 4.8 | 2 | 5.0E-13 | 1.0E-06 |
| CC.12-3 | K417E | 557 | 23 | 4.5 | 1 | 5.0E-13 | 3.1E-13 |
| CC.12-3 | K417F | 344 | 12 | 4.7 | 3 | 5.0E-13 | 6.8E-10 |
| CC.12-3 | K417G | 1439 | 54 | 4.6 | 2 | 5.0E-13 | 1.3E-34 |
| CC.12-3 | K417H | 352 | 17 | 4.2 | 2 | 5.0E-13 | 1.0E-07 |
| CC.12-3 | K417I | 502 | 12 | 5.3 | 1 | 5.0E-13 | 2.3E-17 |
| CC.12-3 | K417L | 2239 | 63 | 5.0 | 2 | 5.0E-13 | 1.2E-65 |
| CC.12-3 | K417M | 983 | 23 | 5.3 | 1 | 5.0E-13 | 5.4E-33 |
| CC.12-3 | K417N | 467 | 19 | 4.5 | 1 | 5.0E-13 | 1.8E-11 |
| CC.12-3 | K417P | 922 | 11 | 6.3 | 2 | 5.0E-13 | 3.5E-40 |
| CC.12-3 | K417Q | 680 | 28 | 4.5 | 1 | 5.0E-13 | 7.1E-16 |
| CC.12-3 | K417R | 1486 | 84 | 4.0 | 1 | 8.7E-13 | 8.3E-23 |
| CC.12-3 | K417S | 1475 | 45 | 4.9 | 2 | 5.0E-13 | 1.5E-41 |

| | | | | | | | |
|---|---|---|---|---|---|---|---|
| CC.12-3 | K417T | 1229 | 38 | 4.9 | 1 | 5.0E-13 | 1.5E-34 |
| CC.12-3 | K417V | 1080 | 33 | 4.9 | 2 | 5.0E-13 | 8.3E-31 |
| CC.12-3 | K417W | 1127 | 42 | 4.6 | 2 | 5.0E-13 | 1.3E-27 |
| CC.12-3 | K417Y | 344 | 15 | 4.4 | 2 | 5.0E-13 | 2.6E-08 |
| CC.12-3 | D420A | 2677 | 84 | 4.9 | 1 | 5.0E-13 | 1.5E-72 |
| CC.12-3 | D420C | 619 | 48 | 3.6 | 2 | 3.0E-07 | 2.5E-06 |
| CC.12-3 | D420E | 2875 | 113 | 4.5 | 1 | 5.0E-13 | 1.0E-64 |
| CC.12-3 | D420K | 317 | 10 | 4.9 | 2 | 5.0E-13 | 7.6E-10 |
| CC.12-3 | D420L | 1313 | 54 | 4.5 | 2 | 5.0E-13 | 3.1E-29 |
| CC.12-3 | D420M | 550 | 26 | 4.3 | 3 | 5.0E-13 | 1.6E-11 |
| CC.12-3 | D420N | 595 | 24 | 4.5 | 1 | 5.0E-13 | 2.5E-14 |
| CC.12-3 | D420Q | 410 | 24 | 4.0 | 2 | 2.6E-12 | 4.6E-07 |
| CC.12-3 | D420R | 2769 | 110 | 4.5 | 2 | 5.0E-13 | 8.6E-62 |
| CC.12-3 | D420S | 1061 | 49 | 4.3 | 2 | 5.0E-13 | 2.4E-21 |
| CC.12-3 | D420T | 530 | 33 | 3.9 | 2 | 4.0E-11 | 5.1E-08 |
| CC.12-3 | D420Y | 273 | 24 | 3.4 | 1 | 2.0E-05 | 7.1E-03 |
| CC.12-3 | Y421A | 1030 | 109 | 3.1 | 2 | 3.1E-03 | 4.8E-04 |
| CC.12-3 | Y421C | 1842 | 126 | 3.7 | 1 | 2.3E-09 | 1.8E-20 |

| | | | | | | | |
|---|---|---|---|---|---|---|---|
| CC.12-3 | Y421F | 729 | 80 | 3.1 | 1 | 7.2E-03 | 6.7E-03 |
| CC.12-3 | Y421H | 881 | 27 | 4.9 | 1 | 5.0E-13 | 2.3E-25 |
| CC.12-3 | Y421L | 3735 | 159 | 4.4 | 2 | 5.0E-13 | 2.6E-77 |
| CC.12-3 | Y421N | 1039 | 56 | 4.1 | 1 | 5.0E-13 | 1.7E-17 |
| CC.12-3 | Y421W | 3746 | 156 | 4.5 | 2 | 5.0E-13 | 2.9E-79 |
| CC.12-3 | N460V | 315 | 28 | 3.4 | 2 | 1.1E-01 | 8.5E-03 |
| CC.12-3 | Q498H | 3844 | 296 | 3.6 | 1 | 2.1E-02 | 4.3E-28 |
| CC.12-3 | Q498W | 4847 | 466 | 3.3 | 2 | 2.1E-01 | 4.7E-17 |
| CC.12-3 | Q498Y | 1445 | 147 | 3.2 | 2 | 3.1E-01 | 4.8E-05 |
| CC.12-3 | N501F | 9266 | 617 | 3.8 | 2 | 2.3E-03 | 7.4E-91 |
| CC.12-3 | N501W | 4284 | 305 | 3.7 | 3 | 6.9E-03 | 1.9E-37 |
| CC.12-3 | N501Y | 9508 | 639 | 3.8 | 1 | 2.7E-03 | 1.9E-91 |
| CC.12-3 | Y505W | 11672 | 894 | 3.6 | 2 | 1.9E-02 | 1.3E-82 |
| CC.12-13 | A475K | 35 | 11 | 3.3 | 2 | 3.1E-06 | 3.7E-02 |
| CC.12-13 | N501W | 2074 | 745 | 3.1 | 3 | 1.2E-04 | 1.3E-35 |
| CC.12-13 | N460P | 21 | 8 | 3.0 | 2 | 5.1E-04 | 1.8E-01 |
| CC.12-13 | E484N | 21 | 8 | 3.0 | 2 | 5.1E-04 | 1.8E-01 |
| CC.12-13 | N501F | 3378 | 1311 | 3.0 | 2 | 7.8E-04 | 3.5E-43 |

| | | | | | | | |
|---|---|---|---|---|---|---|---|
| CC.12-13 | N501V | 4041 | 1593 | 3.0 | 2 | 1.1E-03 | 3.5E-48 |
| CC.12-13 | N501Y | 3028 | 1339 | 2.8 | 1 | 1.2E-02 | 2.4E-21 |
| CC.12-13 | L455M | 19 | 10 | 2.6 | 1 | 2.1E-01 | 4.5E-01 |
| CC.12-13 | N501T | 3053 | 1677 | 2.5 | 1 | 3.6E-01 | 2.1E-03 |
| CC.12-13 | N501I | 2695 | 1531 | 2.4 | 1 | 5.5E-01 | 4.9E-02 |
| CC.6-29 | A475R | 174 | 6 | 4.4 | 2 | 1.1E-07 | 5.6E-04 |
| CC.6-29 | S477P | 472 | 18 | 4.3 | 1 | 1.8E-06 | 7.2E-08 |
| CC.6-29 | T478L | 1562 | 85 | 3.7 | 2 | 3.4E-03 | 2.1E-13 |
| CC.6-29 | T478Q | 239 | 12 | 3.9 | 2 | 8.4E-04 | 1.6E-03 |
| CC.6-29 | T478R | 1421 | 77 | 3.8 | 1 | 3.2E-03 | 1.9E-12 |
| CC.6-29 | E484I | 392 | 25 | 3.5 | 2 | 3.3E-02 | 1.9E-03 |
| CC.6-29 | E484K | 615 | 36 | 3.6 | 1 | 1.1E-02 | 1.9E-05 |
| CC.6-29 | E484R | 1313 | 87 | 3.5 | 2 | 5.2E-02 | 1.5E-07 |
| CC.6-29 | F486A | 741 | 27 | 4.3 | 2 | 5.3E-07 | 4.1E-12 |
| CC.6-29 | F486G | 866 | 35 | 4.2 | 2 | 7.8E-06 | 2.1E-12 |
| CC.6-29 | F486I | 551 | 29 | 3.8 | 1 | 1.9E-03 | 5.6E-06 |
| CC.6-29 | F486L | 2159 | 103 | 3.9 | 1 | 3.1E-04 | 1.4E-22 |
| CC.6-29 | F486R | 325 | 20 | 3.6 | 2 | 2.1E-02 | 3.0E-03 |

| | | | | | | | |
|---|---|---|---|---|---|---|---|
| CC.6-29 | F486S | 1207 | 47 | 4.2 | 1 | 3.1E-06 | 1.9E-17 |
| CC.6-29 | F486V | 1421 | 58 | 4.2 | 1 | 1.0E-05 | 4.0E-19 |
| CC.6-31 | Q493W | 176 | 39 | 3.2 | 2 | 1.8E-05 | 3.3E-04 |
| CC.6-31 | Q493F | 166 | 41 | 3.1 | 3 | 3.0E-04 | 2.9E-03 |
| CC.6-31 | Y505W | 3857 | 1111 | 2.8 | 2 | 8.5E-03 | 1.6E-22 |
| CC.6-31 | L455W | 33 | 10 | 2.8 | 1 | 2.2E-02 | 2.9E-01 |
| CC.6-31 | Q493Y | 124 | 38 | 2.7 | 2 | 2.7E-02 | 9.2E-02 |
| CC.6-31 | N450A | 20 | 8 | 2.4 | 2 | 1.1E+00 | 6.0E-01 |

**Table S3.** Mutations identified in literature. (19–23)

| Mutation | Antibody | Germline | Source(PMID) |
|---|---|---|---|
| T345A | 2H04 | IGHV1-55 | 33535027 |
| T345N | 2H04 | IGHV1-55 | 33535027 |
| T345S | 2H04 | IGHV1-55 | 33535027 |
| R346G | SARS2-01 | | 33535027 |
| R346G | 2H04 | IGHV1-55 | 33535027 |
| R346K | C135 | IGHV3-30 | 32743579 |
| R346M | C135 | IGHV3-30 | 32743579 |
| R346S | C135 | IGHV3-30 | 32743579 |
| A352D | SARS2-01 | | 33535027 |
| Y369C | COV2-2082 | IGHV3-20 | 32935107 |
| N370K | COV2-2082 | IGHV3-20 | 32935107 |
| N370S | COV2-2082 | IGHV3-20 | 32935107 |
| A372S | COV2-2082 | IGHV3-20 | 32935107 |
| A372T | COV2-2082 | IGHV3-20 | 32935107 |
| A372V | COV2-2082 | IGHV3-20 | 32935107 |
| T376I | COV2-2082 | IGHV3-20 | 32935107 |
| T376I | COV2-2094 | IGHV3-20 | 32935107 |

| | | | |
|---|---|---|---|
| **K378E** | **SARS2-31** | | **33535027** |
| **K378N** | **COV2-2677** | **IGHV4-39** | **32935107** |
| **K378N** | **COV2-2082** | **IGHV3-20** | **32935107** |
| **K378N** | **COV2-2094** | **IGHV3-20** | **32935107** |
| **K378Q** | **COV2-2677** | **IGHV4-39** | **33535027** |
| **K378R** | **COV2-2677** | **IGHV4-39** | **32935107** |
| **K378R** | **COV2-2082** | **IGHV3-20** | **32935107** |
| **K378R** | **COV2-2094** | **IGHV3-20** | **32935107** |
| **P384L** | **COV2-2677** | **IGHV4-39** | **32935107** |
| **P384S** | **COV2-2677** | **IGHV4-39** | **32935107** |
| **R408I** | **COV2-2082** | **IGHV3-20** | **32935107** |
| **R408I** | **COV2-2094** | **IGHV3-20** | **32935107** |
| **R408K** | **COV2-2082** | **IGHV3-20** | **32935107** |
| **R408K** | **COV2-2094** | **IGHV3-20** | **32935107** |
| **R408K** | **SARS2-31** | | **33535027** |
| **R408T** | **COV2-2082** | **IGHV3-20** | **32935107** |
| **R408T** | **COV2-2094** | **IGHV3-20** | **32935107** |
| **A411S** | **COV2-2082** | **IGHV3-20** | **32935107** |

| | | | |
|---|---|---|---|
| K417E | REGN10933 | IGHV3-48 | 32540904 |
| K417N | COV2-2082 | IGHV3-20 | 32935107 |
| K417N | COV2-2094 | IGHV3-20 | 32935107 |
| K417R | COV2-2082 | IGHV3-20 | 32935107 |
| K417R | COV2-2094 | IGHV3-20 | 32935107 |
| A435S | COV2-2094 | IGHV3-20 | 32935107 |
| D614G+A435S | H014 | | 32730807 |
| N439K | H00S022 | | 32730807 |
| N440K | C135 | IGHV3-30 | 32743579 |
| L441R | 2H04 | IGHV1-55 | 33535027 |
| K444E | 2H04 | IGHV1-55 | 33535027 |
| K444E | SARS2-38 | | 33535027 |
| K444E | SARS2-22 | | 33535027 |
| K444N | SARS2-38 | | 33535027 |
| K444R | SARS2-22 | | 33535027 |
| K444Q | REGN10934 | IGHV3-15 | 32540904 |
| K444Q | REGN10987 | IGHV3-30 | 32540904 |
| K444R | SARS2-22 | | 33535027 |

| | | | |
|---|---|---|---|
| V445A | REGN10987 | IGHV3-30 | 32540904 |
| V445A | COV2-2499 | IGHV4-39 | 32935107 |
| V445F | COV2-2499 | IGHV4-39 | 32935107 |
| V445G | SARS2-22 | | 33535027 |
| V445I | COV2-2499 | IGHV4-39 | 32935107 |
| G446A | COV2-2096 | IGHV1-8 | 32935107 |
| G446A | COV2-2499 | IGHV4-39 | 32935107 |
| G446D | SARS2-02 | | 33535027 |
| G446D | SARS2-32 | | 33535027 |
| G446D | SARS2-38 | | 33535027 |
| G446D | SARS2-22 | | 33535027 |
| G446S | COV2-2096 | IGHV1-8 | 32935107 |
| G446S | COV2-2499 | IGHV4-39 | 32935107 |
| G446V | COV2-2096 | IGHV1-8 | 32935107 |
| G446V | COV2-2499 | IGHV4-39 | 32935107 |
| G446V | SARS2-02 | | 33535027 |
| N450D | SARS2-07 | | 33535027 |

| | | | |
|---|---|---|---|
| N450K | SARS2-32 | | 33535027 |
| N450Y | SARS2-32 | | 33535027 |
| L452M | COV2-2096 | IGHV1-8 | 32935107 |
| L452R | COV2-2096 | IGHV1-8 | 32935107 |
| L452R | X593 | | 32730807 |
| L452R | P2B-2F6 | IGHV4-38 | 32730807 |
| L452R | SARS2-01 | | 33535027 |
| L452R | SARS2-32 | | 33535027 |
| Y453F | REGN10933 | IGHV3-48 | 32540904 |
| L455F | REGN10933 | IGHV3-48 | 32540904 |
| K458Q | SARS2-66 | | 33535027 |
| I472V | COV2-2479 | IGHV1-69 | 32935107 |
| D614G+I472V | X593 | | 32730807 |
| Q474P | SARS2-34 | | 33535027 |
| A475V | 157 | | 32730807 |
| A475V | 247 | | 32730807 |
| A475V | CB6 | IGHV3-66 | 32730807 |
| A475V | P2C-1F11 | IGHV3-66 | 32730807 |

| | | | |
|---|---|---|---|
| **A475V** | **B38** | **IGHV3-53** | **32730807** |
| **A475V** | **CA1** | **IGHV1-18** | **32730807** |
| **A475V** | **COV2-2165** | **IGHV3-66** | **32935107** |
| **A475V** | **COV2-2832** | **IGHV3-66** | **32935107** |
| **G476D** | **SARS2-21** | | **33535027** |
| **G476D** | **SARS2-34** | | **33535027** |
| **G476D** | **SARS2-71** | | **33535027** |
| **G476S** | **SARS2-21** | | **33535027** |
| **S477G** | **SARS2-16** | | **33535027** |
| **S477G** | **SARS2-07** | | **33535027** |
| **S477G** | **SARS2-19** | | **33535027** |
| **S477G** | **SARS2-34** | | **33535027** |
| **S477G** | **SARS2-58** | | **33535027** |
| **S477G** | **SARS2-71** | | **33535027** |
| **S477I** | **SARS2-58** | | **33535027** |
| **S477N** | **SARS2-16** | | **33535027** |
| **S477N** | **SARS2-07** | | **33535027** |

| | | | |
|---|---|---|---|
| S477N | SARS2-19 | | 33535027 |
| S477N | SARS2-34 | | 33535027 |
| S477N | SARS2-58 | | 33535027 |
| P477L | SARS2-07 | | 33535027 |
| S477R | SARS2-07 | | 33535027 |
| S477R | SARS2-16 | | 33535027 |
| S477R | SARS2-23 | | 33535027 |
| S477R | SARS2-34 | | 33535027 |
| T478I | SARS2-19 | | 33535027 |
| T478I | SARS2-21 | | 33535027 |
| T478I | SASRS2-71 | | 33535027 |
| T478P | SASRS2-71 | | 33535027 |
| P479L | SARS2-34 | | 33535027 |
| P479L | SARS2-71 | | 33535027 |
| P479S | SARS2-21 | | 33535027 |
| V483A | X593 | | 32730807 |
| V483A | P2B-2F6 | IGHV4-38 | 32730807 |

| | | | |
|---|---|---|---|
| V483F | SARS2-23 | | 33535027 |
| V483G | SARS2-23 | | 33535027 |
| E484A | 2B04 | IGHV2-9 | 33535027 |
| E484A | COV2-2832 | IGHV3-66 | 32935107 |
| E484A | COV2-2479 | IGHV1-69 | 32935107 |
| E484A | COV2-2050 | IGHV1-2 | 32935107 |
| E484A | 1B07 | IGHV2-9 | 33535027 |
| E484A | COV2-2096 | IGHV1-8 | 32935107 |
| E484D | COV2-2832 | IGHV3-66 | 32935107 |
| E484D | COV2-2479 | IGHV1-69 | 32935107 |
| E484D | COV2-2050 | IGHV1-2 | 32935107 |
| E484D | COV2-2096 | IGHV1-8 | 32935107 |
| E484D | 1B07 | IGHV2-9 | 33535027 |
| E484D | SARS2-23 | | 33535027 |
| E484D | SARS2-66 | | 33535027 |
| E484G | 1B07 | IGHV2-9 | 33535027 |
| E484K | COV2-2832 | IGHV3-66 | 32935107 |
| E484K | COV2-2479 | IGHV1-69 | 32935107 |

| | | | |
|---|---|---|---|
| **E484K** | **COV2-2050** | **IGHV1-2** | **32935107** |
| **E484K** | **COV2-2096** | **IGHV1-8** | **32935107** |
| **E484K** | **REGN10989** | **IGHV1-2** | **32540904** |
| **E484K** | **REGN10933** | **IGHV3-48** | **32540904** |
| **E484K** | **REGN10934** | **IGHV3-15** | **32540904** |
| **E484K** | **REGN10989/10934** | **IGHV1-2/IGHV3-15** | **32540904** |
| **E484K** | **2B04** | **IGHV2-9** | **33535027** |
| **E484K** | **1B07** | **IGHV2-9** | **33535027** |
| **E484K** | **SARS2-02** | | **33535027** |
| **E484K** | **SARS2-32** | | **33535027** |
| **E484K** | **SARS2-58** | | **33535027** |
| **E484K** | **C121** | **IGHV1-2** | **32743579** |
| **E484K** | **C144** | **IGHV3-53** | **32743579** |
| **E484Q** | **COV2-2832** | **IGHV3-66** | **32935107** |
| **E484Q** | **COV2-2479** | **IGHV1-69** | **32935107** |
| **E484Q** | **COV2-2050** | **IGHV1-2** | **32935107** |
| **E484Q** | **COV2-2096** | **IGHV1-8** | **32935107** |
| **G485D** | **REGN10989** | **IGHV1-2** | **32540904** |

| | | | |
|---|---|---|---|
| **F486S** | **2B04** | **IGHV2-9** | **33535027** |
| **F486S** | **SARS2-21** | | **33535027** |
| **F486V** | **REGN10989** | **IGHV1-2** | **32540904** |
| **F486V** | **REGN10933** | **IGHV3-48** | **32540904** |
| **F486V** | **SARS2-58** | | **33535027** |
| **F486V** | **SARS2-71** | | **33535027** |
| **F486L** | **COV2-2832** | **IGHV3-66** | **32935107** |
| **F486L** | **SARS2-21** | | **33535027** |
| **F486Y** | **1B07** | **IGHV2-9** | **33535027** |
| **F490P** | **REGN10989** | **IGHV1-2** | **32540904** |
| **F490P** | **REGN10934** | **IGHV3-15** | **32540904** |
| **F490P** | **REGN10989/10934** | **IGHV1-2/IGHV3-15** | **32540904** |
| **F490L** | **X593** | | **32730807** |
| **F490L** | **261-262** | | **32730807** |
| **F490L** | **H4** | **IGHV1-2** | **32730807** |
| **F490L** | **P2B-2F6** | **IGHV4-38** | **32730807** |
| **F490L** | **COV2-2479** | **IGHV1-69** | **32935107** |

| | | | |
|---|---|---|---|
| F490L | COV2-2050 | IGHV1-2 | 32935107 |
| F490L | COV2-2096 | IGHV1-8 | 32935107 |
| F490L | SARS2-66 | | 33535027 |
| F490L | C121 | IGHV1-2 | 32743579 |
| F490S | COV2-2479 | IGHV1-69 | 32935107 |
| F490S | COV2-2050 | IGHV1-2 | 32935107 |
| F490S | COV2-2096 | IGHV1-8 | 32935107 |
| F490S | SARS2-32 | | 33535027 |
| Q493K | REGN10989 | IGHV1-2 | 32540904 |
| Q493K | REGN10933 | IGHV3-48 | 32540904 |
| Q493K | REGN10989/10934 | IGHV1-2/IGHV3-15 | 32540904 |
| Q493K | C144 | IGHV3-53 | 32743579 |
| Q493R | C144 | IGHV3-53 | 32743579 |
| Q493K | C121 | IGHV1-2 | 32743579 |
| S494L | COV2-2096 | IGHV1-8 | 32935107 |
| S494P | COV2-2096 | IGHV1-8 | 32935107 |
| S494P | SARS2-01 | | 33535027 |
| P499H | COV2-2499 | IGHV4-39 | 32935107 |

| | | | |
|---|---|---|---|
| **P499R** | **COV2-2499** | **IGHV4-39** | **32935107** |
| **P499S** | **COV2-2499** | **IGHV4-39** | **32935107** |
| **G504D** | **SARS2-31** | | **33535027** |
| **Y508H** | **H014** | | **32730807** |

### Table S4. List of plasmids used in this study.

| Name | Description | *E. coli* Marker | Yeast marker | Source |
|---|---|---|---|---|
| pUC19-S-ecto-B | S ectodomain fragment positions 501-814 with BsaI sites for assembly and BbvCI site | Amp | | This study |
| pUC19-S-ecto-C-Nterm | S ectodomain fragment positions 815-1198 with a C-terminal T4 fibritin trimerization domain with BsaI sites for assembly and BbvCI site | Amp | | This study |
| pUC19-S-ecto-A-Nterm-KanR | S ectodomain fragment positions 13-500 with BsaI sites for assembly and BbvCI site | Kan | | This study |
| pUC19-S-ecto-Nterm | S ectodomain for N-terminal YSD | Kan | | This study |
| pJS697 | YSD vector backbone (C-terminal fusion) for in vivo HR | Kan | TRP1 | This study |
| pJS698 | YSD vector backbone (N-terminal fusion) for in vivo HR | Kan | TRP1 | This study |
| pJS699 | S-RBD(333-537)-N343Q | Kan | | Banach et al., 2021 (4) |

**Table S5. List of primers used.**

| Name | description | sequence |
|---|---|---|
| Downcmyc | Secondary primer for nicking mutagenesis | CAAGTCCTCTTCAGAAATAAGCTTTG |
| M13F | Secondary primer for oligo pool mutagenesis | TGTAAAACGACGGCCAGT |
| MBK-175 | KanR-fwd | CAATAATATTGAAAAAGGAAGAGT |
| MBK-176 | KanR-rev | ATGAGTAAACTTGGTCTGACAGTT |
| MBK-177 | A-pUC19-fwd | AACTGTCAGACCAAGTTTACTCAT |
| MBK-178 | A-pUC19-rev | ACTCTTCCTTTTTCAATATTATTG |
| MBK-180 | DS-tile1-fwd | GTTCAGAGTTCTACAGTCCGACGATCACACGTGGTGTTTATTACCCT |
| MBK-181 | DS-tile1-rev | CCTTGGCACCCGAGAATTCCACATAAGAAAAGGCTGAGAGACATA |
| MBK-301 | DS-tile2-fwd | GTTCAGAGTTCTACAGTCCGACGATCCTTAGGGAATTTGTGTTTAAG |
| MBK-302 | DS-tile2-rev | CCTTGGCACCCGAGAATTCCAAACTTCACCAAAAGGGCACAA |
| MBK-303 | DS-tile3-fwd | GTTCAGAGTTCTACAGTCCGACGATCAGGAAGAGAATCAGCAACTGT |
| MBK-304 | DS-tile3-rev | CCTTGGCACCCGAGAATTCCAATGATTGTAAAGGAAAGTAACA |
| MBK-305 | DS-tile4-fwd | GTTCAGAGTTCTACAGTCCGACGATCAGTAGTAGTACTTTCTTTTGAACTT |
| MBK-306 | DS-tile4-rev | CCTTGGCACCCGAGAATTCCAGGTGTAATGTCAAGAATCTCAAG |
| MBK-307 | DS-tile5-fwd | GTTCAGAGTTCTACAGTCCGACGATCGACTCACTTTCTTCCACAGCA |
| MBK-308 | DS-tile5-rev | CCTTGGCACCCGAGAATTCCAAGCTCTGATTTCTGCAGCTCT |

| | | |
|---|---|---|
| **PJS-P2192** | **pETCON-NK-BsaI-C-term-fwd** | TGTTATGGAGCGGGTCTCAGGGGGCGGATCCGAA |
| **PJS-P2193** | **pETCON-NK-BsaI-C-term-rev** | ACGTTCAGTGATGGTCTCTACTAGCCTGCAGAGC |
| **PJS-P2194** | **pETCON-NK-BsaI-N-term-fwd** | TGTTATGGAGCGGGTCTCACAGGAACTGACAACTATATGC |
| **PJS-P2195** | **pETCON-NK-BsaI-N-term-rev** | ACGTTCAGTGATGGTCTCTGAAAATATTGAAAAACAGCGAAGTAA |
| **RBD_1** | T333-NNK | TGGAGGCGGTAGCGGAGGCGGAGGGTCGNNKAACTTGTGCCCTTTTGGTGAAGTTTTTC |
| **RBD_2** | N334-NNK | AGGCGGTAGCGGAGGCGGAGGGTCGACANNKTTGTGCCCTTTTGGTGAAGTTTTTCAAG |
| **RBD_3** | L335-NNK | CGGTAGCGGAGGCGGAGGGTCGACAAACNNKTGCCCTTTTGGTGAAGTTTTTCAAGCCA |
| **RBD_4** | G339-NNK | CGGAGGGTCGACAAACTTGTGCCCTTTTNNKGAAGTTTTTCAAGCCACCAGATTTGCAT |
| **RBD_5** | E340-NNK | AGGGTCGACAAACTTGTGCCCTTTTGGTNNKGTTTTTCAAGCCACCAGATTTGCATCTG |
| **RBD_6** | A344-NNK | CTTGTGCCCTTTTGGTGAAGTTTTTCAANNKACCAGATTTGCATCTGTTTATGCTTGGA |
| **RBD_7** | T345-NNK | GTGCCCTTTTGGTGAAGTTTTTCAAGCCNNKAGATTTGCATCTGTTTATGCTTGGAACA |
| **RBD_8** | R346-NNK | CCCTTTTGGTGAAGTTTTTCAAGCCACCNNKTTTGCATCTGTTTATGCTTGGAACAGGA |
| **RBD_9** | S349-NNK | TGAAGTTTTTCAAGCCACCAGATTTGCANNKGTTTATGCTTGGAACAGGAAGAGAATCA |
| **RBD_10** | Y351-NNK | TTTTCAAGCCACCAGATTTGCATCTGTTNNKGCTTGGAACAGGAAGAGAATCAGCAACT |
| **RBD_11** | A352-NNK | TCAAGCCACCAGATTTGCATCTGTTTATNNKTGGAACAGGAAGAGAATCAGCAACTGTG |
| **RBD_12** | N354-NNK | CACCAGATTTGCATCTGTTTATGCTTGGNNKAGGAAGAGAATCAGCAACTGTGTTGCTG |
| **RBD_13** | K356-NNK | ATTTGCATCTGTTTATGCTTGGAACAGGNNKAGAATCAGCAACTGTGTTGCTGATTATT |

| | | |
|---|---|---|
| RBD_14 | R357-NNK | TGCATCTGTTTATGCTTGGAACAGGAAGNNKATCAGCAACTGTGTTGCTGATTATTCTG |
| RBD_15 | I358-NNK | ATCTGTTTATGCTTGGAACAGGAAGAGANNKAGCAACTGTGTTGCTGATTATTCTGTCC |
| RBD_16 | S359-NNK | TGTTTATGCTTGGAACAGGAAGAGAATCNNKAACTGTGTTGCTGATTATTCTGTCCTAT |
| RBD_17 | N360-NNK | TTATGCTTGGAACAGGAAGAGAATCAGCNNKTGTGTTGCTGATTATTCTGTCCTATATA |
| RBD_18 | V362-NNK | TTGGAACAGGAAGAGAATCAGCAACTGTNNKGCTGATTATTCTGTCCTATATAATTCCG |
| RBD_19 | A363-NNK | GAACAGGAAGAGAATCAGCAACTGTGTTNNKGATTATTCTGTCCTATATAATTCCGCAT |
| RBD_20 | D364-NNK | CAGGAAGAGAATCAGCAACTGTGTTGCTNNKTATTCTGTCCTATATAATTCCGCATCAT |
| RBD_21 | S366-NNK | GAGAATCAGCAACTGTGTTGCTGATTATNNKGTCCTATATAATTCCGCATCATTTTCCA |
| RBD_22 | V367-NNK | AATCAGCAACTGTGTTGCTGATTATTCTNNKCTATATAATTCCGCATCATTTTCCACTT |
| RBD_23 | N370-NNK | CTGTGTTGCTGATTATTCTGTCCTATATNNKTCCGCATCATTTTCCACTTTTAAGTGTT |
| RBD_24 | A372-NNK | TGCTGATTATTCTGTCCTATATAATTCCNNKTCATTTTCCACTTTTAAGTGTTATGGAG |
| RBD_25 | S373-NNK | TGATTATTCTGTCCTATATAATTCCGCANNKTTTTCCACTTTTAAGTGTTATGGAGTGT |
| RBD_26 | S375-NNK | TTCTGTCCTATATAATTCCGCATCATTTNNKACTTTTAAGTGTTATGGAGTGTCTCCTA |
| RBD_27 | T376-NNK | TGTCCTATATAATTCCGCATCATTTTCCNNKTTTAAGTGTTATGGAGTGTCTCCTACTA |
| RBD_28 | F377-NNK | CCTATATAATTCCGCATCATTTTCCACTNNKAAGTGTTATGGAGTGTCTCCTACTAAAT |
| RBD_29 | K378-NNK | ATATAATTCCGCATCATTTTCCACTTTTNNKTGTTATGGAGTGTCTCCTACTAAATTAA |
| RBD_30 | Y380-NNK | TTCCGCATCATTTTCCACTTTTAAGTGTNNKGGAGTGTCTCCTACTAAATTAAATGATC |
| RBD_31 | G381-NNK | CGCATCATTTTCCACTTTTAAGTGTTATNNKGTGTCTCCTACTAAATTAAATGATCTCT |

| | | |
|---|---|---|
| RBD_32 | V382-NNK | ATCATTTTCCACTTTTAAGTGTTATGGANNKTCTCCTACTAAATTAAATGATCTCTGCT |
| RBD_33 | S383-NNK | ATTTTCCACTTTTAAGTGTTATGGAGTGNNKCCTACTAAATTAAATGATCTCTGCTTTA |
| RBD_34 | P384-NNK | TTCCACTTTTAAGTGTTATGGAGTGTCTNNKACTAAATTAAATGATCTCTGCTTTACTA |
| RBD_35 | T385-NNK | CACTTTTAAGTGTTATGGAGTGTCTCCTNNKAAATTAAATGATCTCTGCTTTACTAATG |
| RBD_36 | K386-NNK | TTTTAAGTGTTATGGAGTGTCTCCTACTNNKTTAAATGATCTCTGCTTTACTAATGTCT |
| RBD_37 | N388-NNK | GTGTTATGGAGTGTCTCCTACTAAATTANNKGATCTCTGCTTTACTAATGTCTATGCAG |
| RBD_38 | D389-NNK | TTATGGAGTGTCTCCTACTAAATTAAATNNKCTCTGCTTTACTAATGTCTATGCAGATT |
| RBD_39 | L390-NNK | TGGAGTGTCTCCTACTAAATTAAATGATNNKTGCTTTACTAATGTCTATGCAGATTCAT |
| RBD_40 | N394-NNK | TACTAAATTAAATGATCTCTGCTTTACTNNKGTCTATGCAGATTCATTTGTAATTAGAG |
| RBD_41 | Y396-NNK | ATTAAATGATCTCTGCTTTACTAATGTCNNKGCAGATTCATTTGTAATTAGAGGTGATG |
| RBD_42 | D405-NNK | CTATGCAGATTCATTTGTAATTAGAGGTNNKGAAGTCAGACAAATCGCTCCAGGGCAAA |
| RBD_43 | R408-NNK | TTCATTTGTAATTAGAGGTGATGAAGTCNNKCAAATCGCTCCAGGGCAAACTGGAAAGA |
| RBD_44 | A411-NNK | AATTAGAGGTGATGAAGTCAGACAAATCNNKCCAGGGCAAACTGGAAAGATTGCTGATT |
| RBD_45 | P412-NNK | TAGAGGTGATGAAGTCAGACAAATCGCTNNKGGGCAAACTGGAAAGATTGCTGATTATA |
| RBD_46 | G413-NNK | AGGTGATGAAGTCAGACAAATCGCTCCANNKCAAACTGGAAAGATTGCTGATTATAATT |
| RBD_47 | Q414-NNK | TGATGAAGTCAGACAAATCGCTCCAGGGNNKACTGGAAAGATTGCTGATTATAATTATA |
| RBD_48 | T415-NNK | TGAAGTCAGACAAATCGCTCCAGGGCAANNKGGAAAGATTGCTGATTATAATTATAAAT |
| RBD_49 | K417-NNK | CAGACAAATCGCTCCAGGGCAAACTGGANNKATTGCTGATTATAATTATAAATTACCAG |

| | | |
|---|---|---|
| **RBD_50** | D420-NNK | CGCTCCAGGGCAAACTGGAAAGATTGCTNNKTATAATTATAAATTACCAGATGATTTTA |
| **RBD_51** | Y421-NNK | TCCAGGGCAAACTGGAAAGATTGCTGATNNKAATTATAAATTACCAGATGATTTTACAG |
| **RBD_52** | K424-NNK | AACTGGAAAGATTGCTGATTATAATTATNNKTTACCAGATGATTTTACAGGCTGCGTTA |
| **RBD_53** | P426-NNK | AAAGATTGCTGATTATAATTATAAATTANNKGATGATTTTACAGGCTGCGTTATAGCTT |
| **RBD_54** | D427-NNK | GATTGCTGATTATAATTATAAATTACCANNKGATTTTACAGGCTGCGTTATAGCTTGGA |
| **RBD_55** | D428-NNK | TGCTGATTATAATTATAAATTACCAGATNNKTTTACAGGCTGCGTTATAGCTTGGAATT |
| **RBD_56** | T430-NNK | TTATAATTATAAATTACCAGATGATTTTNNKGGCTGCGTTATAGCTTGGAATTCTAACA |
| **RBD_57** | N437-NNK | TGATTTTACAGGCTGCGTTATAGCTTGGNNKTCTAACAATCTTGATTCTAAGGTTGGTG |
| **RBD_58** | N439-NNK | TACAGGCTGCGTTATAGCTTGGAATTCTNNKAATCTTGATTCTAAGGTTGGTGGTAATT |
| **RBD_59** | N440-NNK | AGGCTGCGTTATAGCTTGGAATTCTAACNNKCTTGATTCTAAGGTTGGTGGTAATTATA |
| **RBD_60** | L441-NNK | CTGCGTTATAGCTTGGAATTCTAACAATNNKGATTCTAAGGTTGGTGGTAATTATAATT |
| **RBD_61** | K444-NNK | AGCTTGGAATTCTAACAATCTTGATTCTNNKGTTGGTGGTAATTATAATTACCTGTATA |
| **RBD_62** | V445-NNK | TTGGAATTCTAACAATCTTGATTCTAAGNNKGGTGGTAATTATAATTACCTGTATAGAT |
| **RBD_63** | G446-NNK | GAATTCTAACAATCTTGATTCTAAGGTTNNKGGTAATTATAATTACCTGTATAGATTGT |
| **RBD_64** | G447-NNK | TTCTAACAATCTTGATTCTAAGGTTGGTNNKAATTATAATTACCTGTATAGATTGTTTA |
| **RBD_65** | N448-NNK | TAACAATCTTGATTCTAAGGTTGGTGGTNNKTATAATTACCTGTATAGATTGTTTAGGA |
| **RBD_66** | Y449-NNK | CAATCTTGATTCTAAGGTTGGTGGTAATNNKAATTACCTGTATAGATTGTTTAGGAAGT |
| **RBD_67** | N450-NNK | TCTTGATTCTAAGGTTGGTGGTAATTATNNKTACCTGTATAGATTGTTTAGGAAGTCTA |

| | | |
|---|---|---|
| **RBD_68** | L452-NNK | TTCTAAGGTTGGTGGTAATTATAATTACNNKTATAGATTGTTTAGGAAGTCTAATCTCA |
| **RBD_69** | L455-NNK | TGGTGGTAATTATAATTACCTGTATAGANNKTTTAGGAAGTCTAATCTCAAACCTTTTG |
| **RBD_70** | F456-NNK | TGGTAATTATAATTACCTGTATAGATTGNNKAGGAAGTCTAATCTCAAACCTTTTGAGA |
| **RBD_71** | R457-NNK | TAATTATAATTACCTGTATAGATTGTTTNNKAAGTCTAATCTCAAACCTTTTGAGAGAG |
| **RBD_72** | K458-NNK | TTATAATTACCTGTATAGATTGTTTAGGNNKTCTAATCTCAAACCTTTTGAGAGAGATA |
| **RBD_73** | S459-NNK | TAATTACCTGTATAGATTGTTTAGGAAGNNKAATCTCAAACCTTTTGAGAGAGATATTT |
| **RBD_74** | N460-NNK | TTACCTGTATAGATTGTTTAGGAAGTCTNNKCTCAAACCTTTTGAGAGAGATATTTCAA |
| **RBD_75** | K462-NNK | GTATAGATTGTTTAGGAAGTCTAATCTCNNKCCTTTTGAGAGAGATATTTCAACTGAAA |
| **RBD_76** | P463-NNK | TAGATTGTTTAGGAAGTCTAATCTCAAANNKTTTGAGAGAGATATTTCAACTGAAATCT |
| **RBD_77** | F464-NNK | ATTGTTTAGGAAGTCTAATCTCAAACCTNNKGAGAGAGATATTTCAACTGAAATCTATC |
| **RBD_78** | E465-NNK | GTTTAGGAAGTCTAATCTCAAACCTTTTNNKAGAGATATTTCAACTGAAATCTATCAGG |
| **RBD_79** | R466-NNK | TAGGAAGTCTAATCTCAAACCTTTTGAGNNKGATATTTCAACTGAAATCTATCAGGCCG |
| **RBD_80** | I468-NNK | GTCTAATCTCAAACCTTTTGAGAGAGATNNKTCAACTGAAATCTATCAGGCCGGTAGCA |
| **RBD_81** | S469-NNK | TAATCTCAAACCTTTTGAGAGAGATATTNNKACTGAAATCTATCAGGCCGGTAGCACAC |
| **RBD_82** | T470-NNK | TCTCAAACCTTTTGAGAGAGATATTTCANNKGAAATCTATCAGGCCGGTAGCACACCTT |
| **RBD_83** | E471-NNK | CAAACCTTTTGAGAGAGATATTTCAACTNNKATCTATCAGGCCGGTAGCACACCTTGTA |
| **RBD_84** | Y473-NNK | TTTTGAGAGAGATATTTCAACTGAAATCNNKCAGGCCGGTAGCACACCTTGTAATGGTG |
| **RBD_85** | Q474-NNK | TGAGAGAGATATTTCAACTGAAATCTATNNKGCCGGTAGCACACCTTGTAATGGTGTTG |

| | | |
|---|---|---|
| RBD_86 | A475-NNK | GAGAGATATTTCAACTGAAATCTATCAGNNKGGTAGCACACCTTGTAATGGTGTTGAAG |
| RBD_87 | S477-NNK | TATTTCAACTGAAATCTATCAGGCCGGTNNKACACCTTGTAATGGTGTTGAAGGTTTTA |
| RBD_88 | T478-NNK | TTCAACTGAAATCTATCAGGCCGGTAGCNNKCCTTGTAATGGTGTTGAAGGTTTTAATT |
| RBD_89 | P479-NNK | AACTGAAATCTATCAGGCCGGTAGCACANNKTGTAATGGTGTTGAAGGTTTTAATTGTT |
| RBD_90 | N481-NNK | AATCTATCAGGCCGGTAGCACACCTTGTNNKGGTGTTGAAGGTTTTAATTGTTACTTTC |
| RBD_91 | G482-NNK | CTATCAGGCCGGTAGCACACCTTGTAATNNKGTTGAAGGTTTTAATTGTTACTTTCCTT |
| RBD_92 | V483-NNK | TCAGGCCGGTAGCACACCTTGTAATGGTNNKGAAGGTTTTAATTGTTACTTTCCTTTAC |
| RBD_93 | E484-NNK | GGCCGGTAGCACACCTTGTAATGGTGTTNNKGGTTTTAATTGTTACTTTCCTTTACAAT |
| RBD_94 | G485-NNK | CGGTAGCACACCTTGTAATGGTGTTGAANNKTTTAATTGTTACTTTCCTTTACAATCAT |
| RBD_95 | F486-NNK | TAGCACACCTTGTAATGGTGTTGAAGGTNNKAATTGTTACTTTCCTTTACAATCATATG |
| RBD_96 | N487-NNK | CACACCTTGTAATGGTGTTGAAGGTTTTNNKTGTTACTTTCCTTTACAATCATATGGTT |
| RBD_97 | Y489-NNK | TTGTAATGGTGTTGAAGGTTTTAATTGTNNKTTTCCTTTACAATCATATGGTTTCCAAC |
| RBD_98 | F490-NNK | TAATGGTGTTGAAGGTTTTAATTGTTACNNKCCTTTACAATCATATGGTTTCCAACCCA |
| RBD_99 | L492-NNK | TGTTGAAGGTTTTAATTGTTACTTTCCTNNKCAATCATATGGTTTCCAACCCACTAATG |
| RBD_100 | Q493-NNK | TGAAGGTTTTAATTGTTACTTTCCTTTANNKTCATATGGTTTCCAACCCACTAATGGTG |
| RBD_101 | S494-NNK | AGGTTTTAATTGTTACTTTCCTTTACAANNKTATGGTTTCCAACCCACTAATGGTGTTG |
| RBD_102 | Q498-NNK | TTACTTTCCTTTACAATCATATGGTTTCNNKCCCACTAATGGTGTTGGTTACCAACCAT |
| RBD_103 | P499-NNK | CTTTCCTTTACAATCATATGGTTTCCAANNKACTAATGGTGTTGGTTACCAACCATACA |

| | | |
|---|---|---|
| **RBD_104** | T500-NNK | TCCTTTACAATCATATGGTTTCCAACCCNNKAATGGTGTTGGTTACCAACCATACAGAG |
| **RBD_105** | N501-NNK | TTTACAATCATATGGTTTCCAACCCACTNNKGGTGTTGGTTACCAACCATACAGAGTAG |
| **RBD_106** | V503-NNK | ATCATATGGTTTCCAACCCACTAATGGTNNKGGTTACCAACCATACAGAGTAGTAGTAC |
| **RBD_107** | G504-NNK | ATATGGTTTCCAACCCACTAATGGTGTTNNKTACCAACCATACAGAGTAGTAGTACTTT |
| **RBD_108** | Y505-NNK | TGGTTTCCAACCCACTAATGGTGTTGGTNNKCAACCATACAGAGTAGTAGTACTTTCTT |
| **RBD_109** | Q506-NNK | TTTCCAACCCACTAATGGTGTTGGTTACNNKCCATACAGAGTAGTAGTACTTTCTTTTG |
| **RBD_110** | Y508-NNK | ACCCACTAATGGTGTTGGTTACCAACCANNKAGAGTAGTAGTACTTTCTTTTGAACTTC |
| **RBD_111** | E516-NNK | ACCATACAGAGTAGTAGTACTTTCTTTTNNKCTTCTACATGCACCAGCAACTGTTTGTG |
| **RBD_112** | L517-NNK | ATACAGAGTAGTAGTACTTTCTTTTGAANNKCTACATGCACCAGCAACTGTTTGTGGAC |
| **RBD_113** | L518-NNK | CAGAGTAGTAGTACTTTCTTTTGAACTTNNKCATGCACCAGCAACTGTTTGTGGACCTA |
| **RBD_114** | H519-NNK | AGTAGTAGTACTTTCTTTTGAACTTCTANNKGCACCAGCAACTGTTTGTGGACCTAAAA |
| **RBD_115** | A520-NNK | AGTAGTACTTTCTTTTGAACTTCTACATNNKCCAGCAACTGTTTGTGGACCTAAAAAGT |
| **RBD_116** | P521-NNK | AGTACTTTCTTTTGAACTTCTACATGCANNKGCAACTGTTTGTGGACCTAAAAAGTCTA |
| **RBD_117** | A522-NNK | ACTTTCTTTTGAACTTCTACATGCACCANNKACTGTTTGTGGACCTAAAAAGTCTACTA |
| **RBD_118** | T523-NNK | TTCTTTTGAACTTCTACATGCACCAGCANNKGTTTGTGGACCTAAAAAGTCTACTAATT |
| **RBD_119** | P527-NNK | TCTACATGCACCAGCAACTGTTTGTGGANNKAAAAAGTCTACTAATTTGGTTAAAAACA |
| **IFU-104** | L1_Inner_FWD | gttcagagttctacagtccgacgatcTGGAGGAGGCTCTGG |
| **IFU-105** | L1_Inner_REV | ccttggcacccgagaattccaCCAAGCTATAACGCAGCC |

| | | |
|---|---|---|
| IFU-106 | L2_Inner_FWD | gttcagagttctacagtccgacgatcGGCTGCGTTATAGCTTGG |
| IFU-107 | L2_Inner_REV | ccttggcacccgagaattccaGCCCCCTTTGTTTTTAACCAA |
| | Forward Outer Primer | AATGATACGGCGACCACCGAGATCTACACGTTCAGAGTTCTACAGTCCGACGATC |
| | Reverse Outer Primer | CAAGCAGAAGACGGCATACGAGATNNNNNNGTGACTGGAGTTCCTTGGCACCCGAGAATTCCA<br><br>Where NNNNNN is the indexing barcode, for full sequences refer to (24) |

**Data S1.**
Processed files for all nAb escape mutant identification using S RBD.

**Data S2.**
Spike mutational designs and DNA sequences encoded in the oligo pool library.

**Data S3.**
Processed Spike ectodomain library results for sorting against ACE2, CR3022.
